# Sialylated CD43 is a glyco-immune checkpoint for macrophage phagocytosis

**DOI:** 10.1101/2025.05.05.652090

**Authors:** Jooho Chung, Mounica Vallurupalli, Sarah Noel, Gail Schor, YuhJong Liu, Celeste Nobrega, Jonathan J. Perera, Ewa Wrona, Margaret Hu, Yunkang Lin, David W. Wu, Maria Saberi, Ilario Scapozza, Aidan Cruickshank, Elliot C. Woods, Cun Lan Chuong, Filippo Birocchi, Ashwin V. Kammula, Omar I. Avila, Mustafa Kocak, John G. Doench, Dean Procter, Lindsey Thornton, Andrew M. Brunner, Eric Winer, Daniel J. DeAngelo, Jacqueline S. Garcia, Richard M. Stone, Russell W. Jenkins, Marcela V. Maus, Timothy A. Graubert, Kathleen B. Yates, Todd R. Golub, Robert T. Manguso

**Affiliations:** Broad Institute of Harvard and Massachusetts Institute of Technology; Cambridge, MA USA; Krantz Family Center for Cancer Research, Massachusetts General Hospital, Harvard Medical School; Charlestown, MA, USA; Division of Hematology and Oncology, Massachusetts General Brigham Cancer Center; Boston, MA USA; Department of Medical Oncology, Dana-Farber Cancer Institute; Boston, MA USA; Department of Pediatric Oncology, Dana-Farber Cancer Institute, Harvard Medical School; Boston, MA, USA; Department of Clinical and Laboratory Genetics, Medical University of Lodz; Lodz, Poland; Department of Chemotherapy, Copernicus Memorial Hospital; Lodz, Poland

## Abstract

Macrophages in the tumor microenvironment exert potent anti-tumorigenic activity through phagocytosis. Yet therapeutics that enhance macrophage phagocytosis have not improved outcomes in clinical trials for patients with acute myeloid leukemia (AML) or myelodysplastic syndrome (MDS). To systematically identify regulators of phagocytosis, we performed genome-scale CRISPR knockout screens in human leukemia cells co-cultured with human monocyte-derived macrophages. Surprisingly, we found that whereas the classic “don’t eat me” signal CD47 inhibited mouse macrophages, it did not inhibit phagocytosis by human macrophages. In contrast, the O-linked glycosylation and sialylation pathways were strong negative regulators of phagocytosis. In AML, the cell surface O-linked glycoprotein CD43 was the major effector of the O-linked glycosylation and sialylation pathways. Genetic deletion or antibody blockade of CD43 enhanced macrophage phagocytosis. This work highlights the importance of using human platforms to identify immune checkpoints, and nominates CD43 as a glyco-immune regulator of human macrophage phagocytosis.

## Main Text

Phagocytosis is a core function of macrophages that promotes tumor clearance and enhances adaptive immunity through antigen processing and cross-presentation to T lymphocytes. Macrophages phagocytose tumor cells through Fc-receptor mediated recognition of antibody-opsonized targets (antibody-dependent cellular phagocytosis, ADCP) and/or antibody-independent phagocytosis (AICP), which involves the integration of pro-phagocytic “eat me” signals and anti-phagocytic “don’t eat me” signals (*1*). Preclinical studies have identified CD47 on tumor cells as a major “don’t eat me” signal that transmits inhibitory signals to macrophages to restrain phagocytosis (*2*, *3*). However, recent phase III clinical trials of CD47-neutralizing antibodies for the treatment of MDS and AML have shown limited clinical efficacy (*4*). The mechanism explaining these clinical results is unknown, and highlights the need for the discovery of new therapeutic targets with potential to activate the phagocytosis of cancer cells.

To define the complete landscape of tumor-intrinsic macrophage inhibitory checkpoints, we performed genome-scale CRISPR knockout screens of human AML cell lines co-cultured with human macrophages in the presence or absence of antibody opsonization. This approach allowed us to catalog and validate the genes that impact human macrophage-mediated AICP and ADCP. As there are notable differences in receptor expression, activation, and function between mouse and human macrophages (*5*, *6*), we used healthy human donor-derived monocytes differentiated into macrophages to perform unbiased genome-scale screens. We focused on AML because it is refractory to T-cell directed immune checkpoint blockade (*7*, *8*) and has a tumor microenvironment that is rich in myeloid cells (*9*). Here, we show that CD47 is only a weak regulator of human macrophage phagocytosis, whereas the O-linked sialoglycoprotein CD43 is a potent inhibitor of human macrophage AICP and ADCP.

## Results

### Development of a human macrophage co-culture assay to determine regulators of phagocytosis

To establish a scalable and sensitive phagocytosis assay for unbiased genetic screens, we isolated monocytes from healthy donors and evaluated the mode of monocyte differentiation (M-CSF vs GM-CSF), macrophage stimulation (no stimulation, IL-6, IFNγ, TNFα, or LPS), and effector to target ratios on the phagocytosis of two human leukemia cell lines, MV411 and MOLM13 (*10*). Phagocytosis was quantified with flow cytometry to measure the absolute number of remaining leukemia cells after 18-24 hours of co-culture compared to leukemia cells grown in the absence of macrophages (**Fig. S1A**). As a complementary approach, we evaluated phagocytosis of pHrodo-labeled leukemia cells with live-cell imaging, which allowed us to identify macrophages that had recently ingested leukemia cells (**Fig. S1B)**.

Baseline phagocytosis of leukemia cells by unstimulated human macrophages was low (**Fig. S1B-C**). IFNγ stimulation had the most potent effect on macrophage phagocytosis, enhancing absolute phagocytosis by 4-6 fold (**Fig. S1B-C**). M-CSF differentiated macrophages exhibited increased phagocytosis compared to GM-CSF differentiated macrophages at all effector:target ratios (**Fig. S1D**) and this mode of differentiation was selected for further phagocytosis assays.

### Genome-scale CRISPR screens reveal pathways that regulate antibody-independent tumor phagocytosis

To define the tumor-intrinsic factors that regulate AICP, we performed pooled genome-scale CRISPR knockout screens by co-culturing CRISPR-edited leukemia cells with IFNγ-stimulated human macrophages (**Fig. 1A**). To increase generalizability, we performed the screen in two human AML cell lines (MV411 and MOLM13), using different CRISPR enzymes and genome-targeting libraries (Cas9 and Cas12a, respectively). After 18 hours of co-culture, the pool of remaining CRISPR-edited leukemia cells were separated from the adherent macrophages, allowing us to sequence the single guide RNA (sgRNA) barcodes from both the leukemia and macrophage fractions (**Fig. 1A**). The leukemia fraction contained barcoded cells that had escaped phagocytosis while the macrophage fraction contained barcodes derived from recently phagocytosed cells. Quality control parameters demonstrated excellent screen performance, with the majority of sgRNAs being well represented in all conditions (**Fig. S2A-B**). Guide RNA representation in each replicate was highly correlated to the original library (Pearson correlation > 0.8 in leukemia fraction and macrophage fractions) (**Fig. S2C-F**).

**Fig. 1:**
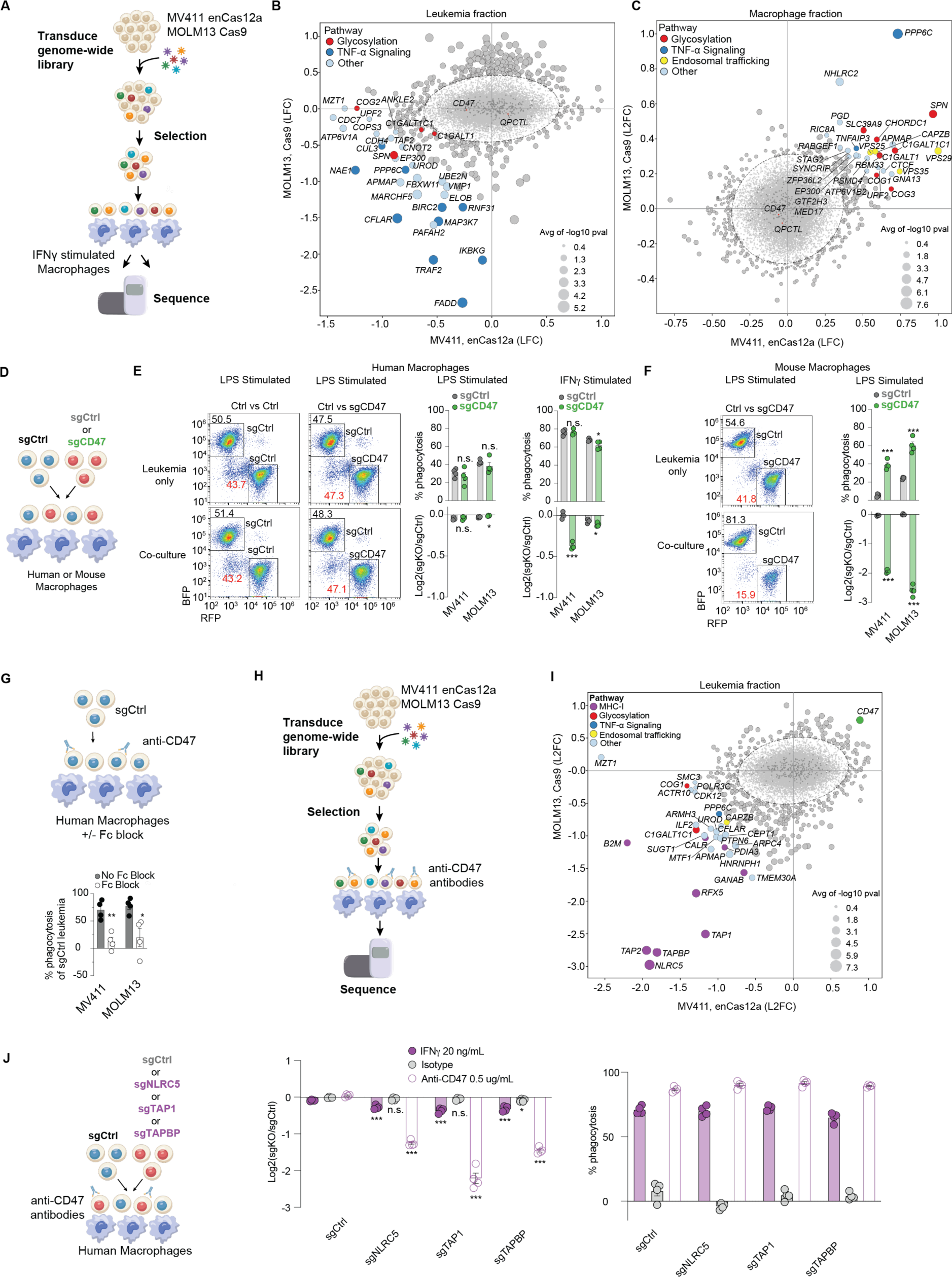
Genome-wide CRISPR screens identify key pathways that regulate antibody-dependent cellular phagocytosis and antibody-independent phagocytosis of human leukemia. **a,** Schematic of antibody-independent cellular phagocytosis screen design. **b,** Scatter plot of log fold change of enrichment or depletion of genes in the leukemia fraction of MV411 (*x*-axis) or MOLM13 (*y*-axis) co-cultured with macrophages versus leukemia alone. Circle size indicates average -log10 (p-value) across both cell lines. **c,** Scatter plot of log fold change of enrichment or depletion of genes in the macrophage fraction of MV411 (x-axis) or MOLM13 (y-axis) co-cultured with macrophages versus leukemia alone. Circle size indicates average -log10 (p-value) across both cell lines. **d**, Schematic of competitive co-culture experiment of human leukemia cells with human or mouse macrophages. **e,** Flow cytometry plot of leukemia cells with and without CD47 knockout cultured alone or with human macrophages stimulated with LPS. Barplots show quantification of the percent of total cells phagocytosed and log2 normalized ratio of knockout cells versus control. Data show the mean ± s.e.m. of four technical replicates and are representative of two different experiments. **f,** Flow cytometry plot of human leukemia cells co-cultured with LPS stimulated mouse macrophages. Barplots shown quantify the percent of phagocytosed cells and ratio of knockout to control remaining after macrophage co-culture. Data show the mean ± s.e.m. of four technical replicates and are representative of two different experiments. **g,** Schematic of co-culture experiment of human leukemia treated with anti-CD47 (MIAP410) antibody prior to co-culture with human macrophages pre-treated with or without Fc receptor blocking antibodies. Barplot shows the percent of phagocytosed cells in the presence or absence of macrophage Fc receptor blockade. Data show the mean ± s.e.m. of four technical replicates and are representative of two different experiments. **h,** Schematic of antibody-dependent cellular phagocytosis screen design. **i,** Scatter plot of log fold change of enrichment or depletion of genes in the leukemia fraction of anti-CD47 treated MV411 (*x*-axis) or MOLM13 (*y*-axis) co-cultured with macrophages versus leukemia alone. Circle size indicates average log10 (p-value) across both cell lines. **j,** Competitive co-culture experiment of control or knockout leukemia cells co-cultured with human macrophages. Log2 normalized ratio of the knockout to control and percent of phagocytosed leukemia are plotted. Data in **e-g, j** were analyzed by unpaired, two-sided Student’s *t*-test, * p<0.05, ** p<0.01, *** p<0.001.

We compared the sgRNA abundances in the co-cultured leukemia and macrophage fractions to leukemia cells cultured alone (**Fig. 1B-C**, **Fig. S3A-D, Table S1-S4**). Consistent with previous reports, genetic deletion of *APMAP* in MV411 or MOLM13 cells resulted in depletion in the leukemia fraction and enrichment in the macrophage fraction, giving us confidence that our co-culture system can identify known regulators of macrophage phagocytosis (**Fig. 1B-C, Fig. S3A-B**) (*11*). Integrated analyses of leukemic and macrophage fractions in each screen revealed additional genes that scored as shared negative regulators of macrophage phagocytosis, including genes involved in O-linked glycosylation (*SPN*, *C1GALT1*, *C1GALT1C1*), TNF alpha signaling (*PPP6C*), and endosomal trafficking (*VPS25*), as well as genes that scored as cell line-specific regulators, including *CDC7, ATP6V1A*, *MZT1* and *VPS29* in MV411, and *FADD*, *IKBKG*, *MAP3K7* and *NHLRC2* in MOLM13 (**Fig. S3A-D)**.

### Genetic loss of CD47 in human leukemia does not potently enhance phagocytosis by human macrophages

The “don’t eat me” signal CD47 is upregulated in AML cells and has been reported to promote evasion of macrophage phagocytosis through its cognate inhibitory receptor, SIRPα (*2*). Anti-CD47 antibodies enhance phagocytosis in leukemia xenograft models in NOD SCID *IL2rg^-/^*^-^ (NSG) mice, providing motivation for testing anti-CD47 antibodies in clinical trials (*3*). We confirmed that treatment of leukemia cells with anti-CD47 antibody (clone MIAP410) potently enhances phagocytosis by unstimulated macrophages (**Fig. S4A**). In contrast, genetic deletion of *CD47* in leukemia cells did not result in depletion in the leukemia fraction or enrichment in the macrophage fraction in our AICP screens (**Fig. 1B-C, Fig. S3A-D**). Similarly, genetic deletion of *QPCTL*, a CD47-modifying enzyme required for optimal binding of CD47 to SIRPα (*12*), did not enhance phagocytosis (**Fig. 1B-C, Table S1-S4**). To ensure that these findings did not represent a false-negative result, we deleted *CD47* in the MV411 and MOLM13 cell lines (**Fig. S4B-C**) and evaluated the functional impact of CD47 loss in competitive phagocytosis experiments by co-culturing 1:1 mixes of control and CD47-deficient leukemias with M-CSF differentiated human macrophages stimulated with LPS or IFNγ (**Fig. 1D**). Consistent with our screen results, *CD47* deletion did not enhance the phagocytosis of human leukemia cells relative to control leukemias by human macrophages (**Fig. 1E**). Similarly, addition of soluble recombinant SIRPα, which antagonizes binding of CD47 on leukemia cells to SIRPα on human macrophages, did not enhance phagocytosis (**Fig. S4D**). To determine whether this represents a difference in the activity of this checkpoint between species, we co-cultured CD47-deficient human leukemias with M-CSF differentiated mouse bone marrow-derived macrophages and observed a significant enhancement of phagocytosis (**Fig. 1F**). Therefore, CD47 expression in human AML cells potently inhibits phagocytosis by mouse but not human macrophages.

Given that genetic deletion of CD47 had a marginal impact on phagocytosis by human macrophages, we assessed whether the mechanism by which anti-CD47 antibodies enhance phagocytosis is mediated through recruitment of FcγR and subsequent antibody-dependent cellular phagocytosis (ADCP). Leukemia cells treated with anti-CD47 antibody (MIAP410) exhibited dramatically enhanced phagocytosis when co-cultured with human macrophages (**Fig. 1G**, grey bars). However, when macrophages were pretreated with an Fc receptor-blocking cocktail of antibodies (anti-CD16/CD32/CD64), there was a significant decrease in the percent of phagocytosed cells, suggesting that antibodies targeting human CD47 exert pro-phagocytic effects through ADCP (**Fig. 1G**, white bars). These findings suggest that anti-CD47 antibodies enhance phagocytosis primarily via antibody-dependent recruitment of macrophage FcγRs, rather than CD47 serving as a “don’t eat me” macrophage inhibitory checkpoint. These results are also consistent with recent work highlighting the importance of Fc-domain optimization to achieve maximal efficacy of anti-CD47 antibodies (*13*, *14*).

### MHC class I potently inhibits antibody-dependent cellular phagocytosis

As tumor antigen-directed antibodies can elicit phagocytosis (*15*, *16*), we next used our screening approach to identify regulators of ADCP (**Fig. 1H**). Given the high expression of CD47 on leukemia cell lines and our findings that genetic loss of CD47 does not impact human AICP (**Fig. 1B-E**), we used an anti-CD47 antibody (clone MIAP410) with strong FcγR activity as an opsonizing antibody to elicit ADCP. We again observed excellent screen performance with good sgRNA representation across all conditions and replicates (Pearson correlation > 0.8 in the leukemia fraction and >0.7 in the macrophage fraction) (**Fig. S5A-F**). Loss of CD47 in leukemia cells was the strongest resistance factor in this screen, consistent with immune escape due to antigen loss (**Fig. 1I**). In contrast to prior cross-species macrophage co-culture screens (*11*), we identified several genes involved in MHC class I expression (*NLRC5*, *TAP1*, *TAP2*, *TAPBP*, *RFX5*, and *B2M*) as the top negative regulators of ADCP in leukemia cells co-cultured with macrophages (**Fig. 1I, Fig. S6, Table S5,S6).** Competition assays confirmed that genetic deletion of *NLRC5*, *TAP1*, or *TAPBP* in MV411 leukemia cells resulted in increased phagocytosis in co-culture with human macrophages stimulated with anti-CD47 antibody but not with IFNγ (**Fig. 1j**), demonstrating that MHC-I expression on AML cells is a major regulator of ADCP but not AICP. We observed similar enhancement of phagocytosis upon treatment of leukemia cells with an opsonizing antibody targeting CD33, another highly expressed leukemia cell surface receptor (**Fig. S7A**).

MHC-I proteins interact with the myeloid-specific inhibitory receptors *LILRB1* and *LILRB2,* which contain immunoreceptor tyrosine-based inhibitory motifs that restrain immune signaling (*17–20*). Mouse macrophages do not express the *LILRB* family of genes and instead express the murine ortholog, *Pirb*(*21, 22*). This species difference in receptor expression highlights the failure of cross-species co-culture screens to resolve the MHC-I::LILRB1 interaction as a key regulator of ADCP(*11*). In human macrophages, genetic loss of *LILRB1* or antibody-mediated blockade of *LILRB1* has been shown to enhance ADCP (*23*, *24*). As there are several clinical trials exploring strategies to disrupt the MHC-I:LILRB axis to enhance immunotherapy response (*25*, *26*), we sought to assess whether genetic loss of MHC-I in leukemia cells enhances ADCP through loss of interaction with *LILRB1* and/or another member of the *LILRB* family, *LILRB2.* We used CRISPR/Cas9 to delete *LILRB1* or *LILRB2* in human macrophages and found that deletion of either gene reduced the preferential phagocytosis of MHC-I-deficient leukemias by macrophages (**Fig. S7B-C**), demonstrating that MHC-I expression can restrain ADCP of AML cells through not just LILRB1 but also LILRB2 on human macrophages.

### Mucin-type O-linked glycosylation is a negative regulator of AICP and ADCP

To systematically rank the impact of individual genes on AICP or ADCP, we integrated data from both the leukemia and macrophage fractions of each screen across MV411 and MOLM13 cell lines. For each individual gene, we calculated an aggregate score of sgRNA depletion in the leukemia fraction and sgRNA enrichment in the macrophage fraction. We found that *PPP6C, SPN, APMAP*, *VPS29,* and *NHLRC2* were the top negative regulators of AICP (**Fig. 2A**). *NLRC5, TAPBP, TAP2, PPP6C,* and *PTPN6* were the top negative regulators of ADCP (**Fig. 2A**). To validate our hit calling method, we showed that genetic deletion of PTPN6 in leukemia cells strongly enhances ADCP in co-culture competition assays (**Fig. 2B, Fig. S7D**).

**Fig. 2:**
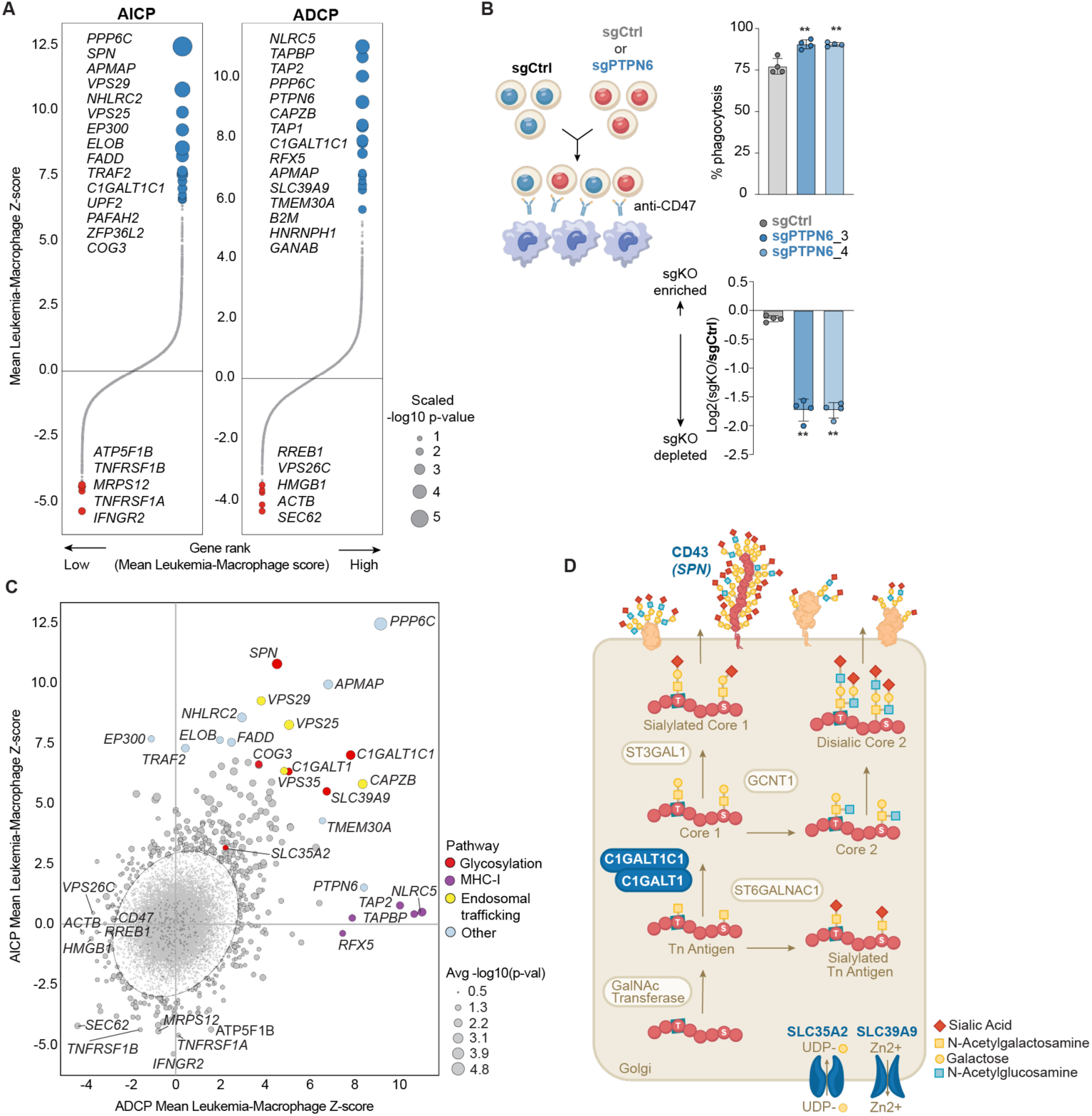
Integrated analysis of genome-scale antibody-dependent and antibody-independent cellular phagocytosis screens reveals O-glycan pathway genes as key regulators of macrophage phagocytosis. **a**, Genes ranked by the average of the difference between the log fold change of enrichment or depletion of the leukemia fraction and macrophage fraction. Circle size is scaled -log10 (p-value). Top 15 enriched genes (blue) and top 5 depleted genes (red) are listed and arranged by statistical significance. **b,** Schematic of competitive co-culture experiment of PTPN6 knockout leukemia. Percent of phagocytosed cells and log2 ratio of knockout to control are shown. Data represent mean ± s.e.m. of four technical replicates and are representative of two independent experiments. **c,** Scatter plot of the difference between the log fold change of enrichment or depletion in the leukemia fraction and macrophage fraction in the ADCP screen (*x*-axis) and AICP screen (*y*-axis). Circle size is the average -log10(p-value). Colors denote pathway annotation. **d,** Schematic of the O-glycosylation pathway highlighting top gene knockouts that enhance AICP and ADCP (blue text). Data in **b** were analyzed by unpaired, two-sided Student’s *t*-test, ** p< 0.01

Next, we focused on hits that regulate both AICP and ADCP by plotting the z-normalized aggregated sgRNA scores in the ADCP and AICP screens (**Fig. 2C**). This approach highlighted *APMAP*, a previously known negative regulator of ADCP(*11*), as a regulator of both AICP and ADCP. Additionally, deletion of *TMEM30A*, which encodes the phosphatidylserine flippase CDC50A, enhanced overall macrophage phagocytosis, likely due to an increase in surface phosphatidylserine exposure which activates scavenger receptors such as MERTK (*27–29*). Most strikingly, several genes in the O-linked glycosylation pathway, including *C1GALT1, C1GALT1C1*, *SLC39A9*, and *SLC35A2*, emerged as major negative regulators of both AICP and ADCP (**Fig. 2C-D**). *SPN,* encoding CD43, a cell surface glycoprotein that undergoes extensive O-glycosylation, also scored as a negative regulator of both AICP and ADCP (**Fig. 2C-D**), implicating the O-glycosylation pathway as a major inhibitor of macrophage phagocytosis. O-linked glycosylation is a post-translational modification of cell surface and secreted proteins that attaches glycan chains terminating in the monosaccharide sialic acid (**Fig. 2D**). C1GALT1 encodes the T-synthase enzyme, which catalyzes an early step in the O-glycosylation pathway, while C1GALT1C1 is a chaperone protein required for C1GALT1 stability (*30*). SLC39A9, a putative zinc transporter, regulates C1GALT1 expression (*31*), and SLC35A2 transports the donor substrate UDP-galactose, which is incorporated into O-glycan structures.

### O-linked glycosylation inhibits phagocytosis through terminal sialylation of cell surface glycoproteins

We generated single gene knockouts of *C1GALT1* and *C1GALT1C1* and evaluated changes in cell surface glycosylation with fluorescently-labeled lectins, which bind unique carbohydrate motifs with high specificity. As expected, *C1GALT1*- and *C1GALT1C1*-deficient leukemias expressed increased levels of the C1GALT1 substrate Tn antigen, decreased levels of the downstream enzymatic product T antigen, and decreased levels of sialic acid (**Fig. S8A**). Knockout of neither *C1GALT1* nor *C1GALT1C1* altered cell growth kinetics *in vitro* (**Fig. S8B**). However, upon co-culture with macrophages, *C1GALT1*- and *C1GALT1C1*-deficient leukemias were more sensitive to both AICP and ADCP (**Fig. 3A, Fig. S8C**). Similarly, genetic deletion of *SLC39A9* or *SLC35A2* also enhanced phagocytosis by IFNγ-stimulated macrophages in competition assays (**Fig. 3B**). In contrast, CRISPR-mediated genetic deletion of *MGAT1*, a key component of the N-linked glycosylation pathway, had no impact on AICP (**Fig. S8D-E**). These data implicate mucin-type O-linked glycosylation, but not N-linked glycosylation, as a major inhibitory pathway for human macrophage phagocytosis.

**Fig. 3:**
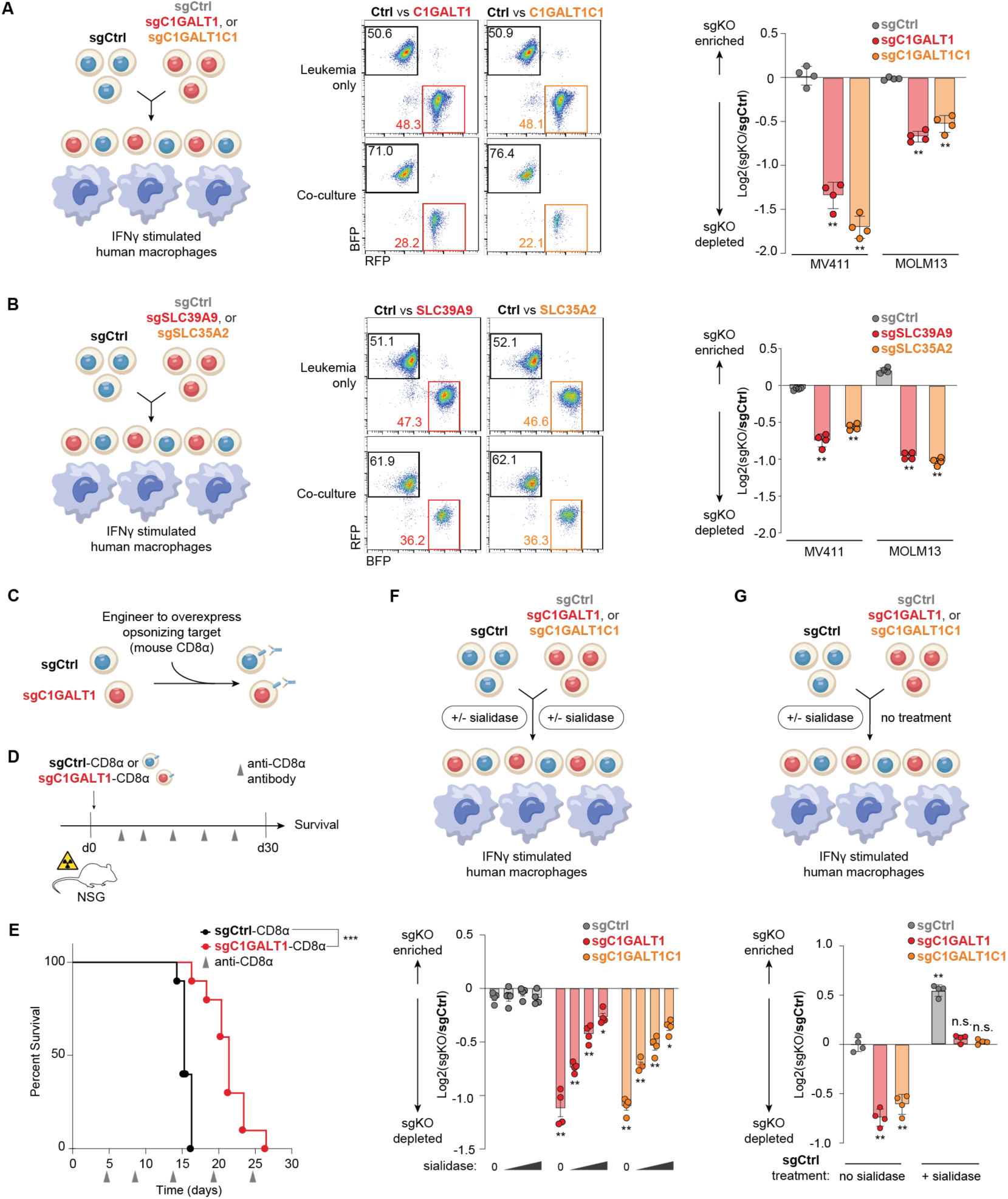
The O-linked glycosylation pathway inhibits human macrophage phagocytosis of leukemia cells through terminal sialic acid residues **a,** Schematic of competitive phagocytosis assay, representative flow cytometry plots, and bar graphs of relative phagocytosis of *C1GALT1*-deficient*, C1GALT1C1-*deficient, or control versus control leukemia cells (MV411 and MOLM13). **b,** Schematic of competitive phagocytosis assay, representative flow cytometry plots, and bar graphs of relative phagocytosis of *SLC39A9-*deficient*, SLC35A2-*deficient, or control versus control sgRNA leukemia cells (MV411 and MOLM13) **c.** Antibody-dependent cellular phagocytosis (ADCP) strategy: control or *C1GALT1*-deficient MV411 cells were engineered to ectopically express mouse CD8α **d.** *In vivo* ADCP model: control or *C1GALT1*-deficient were intravenously engrafted into sublethally irradiated NOD SCID *IL2rg^-/^*^-^ mice. Mice bearing *C1GALT1*-deficient or control MV411 cells were intraperitoneally injected with systemic anti-CD8α antibodies starting on day +5. **e.** Survival of mice challenged with *C1GALT1*-deficient or control MV411 leukemia cells. **f,** Schematic of competitive phagocytosis assay experimental design and bar graphs of relative phagocytosis of *C1GALT1, C1GALT1C1,* or control sgRNA versus control sgRNA leukemia cells pre-treated with varying doses of *V. cholerae* sialidase prior to phagocytosis assays. Both knockout and control cells were pre-treated with sialidase prior to co-culture. **g**, Schematic of competitive phagocytosis assay and bar graphs of relative phagocytosis of *C1GALT1-*deficient*, C1GALT1C1-*deficient, or control MV411 leukemia cells versus control cells pre-treated with *V. cholerae* sialidase prior to phagocytosis assays. Only control cells (but not knockout cells) were pre-treated with sialidase prior to co-culture. For **a-b** and **f-g**, genetically modified or sialidase-treated leukemia cells were co-cultured with or without IFNγ-stimulated macrophages for 18-24 hours. Bar graphs indicate log (fold change) of the ratio of knockout cells relative to control after co-culture with macrophages. Data represent mean ± s.d. of four technical replicates and are representative of 3-5 independent experiments For **e**, data represent three independent survival experiments.

To assess the impact of O-linked glycosylation on macrophage phagocytosis *in vivo*, we established a model of ADCP for both *in vitro* and *in vivo* studies. We decoupled ADCP from confounding effects of interfering with the function of endogenous leukemia antigens by engineering MV411 leukemia cells to ectopically express mouse CD8α (**Fig. 3C, Fig. S8F)**. In competition assays, ectopic expression of mouse CD8α in control leukemias (sgCtrl-CD8α OE) rendered them sensitive to anti-mouse CD8α-mediated ADCP when mixed with control leukemias that overexpressed luciferase (sgCtrl-Luciferase OE) (**Fig. S8G**). *C1GALT1*-deficient leukemias overexpressing CD8α (sgC1GALT1-CD8α), but not *CD47*-deficient leukemias overexpressing CD8α, were selectively phagocytosed compared to sgCtrl-CD8α OE leukemias (**Fig. S8H)**. Thus, ectopic expression of mouse CD8α on MV411 leukemias enables anti-CD8α-mediated cellular phagocytosis and preserves the inhibitory impact of O-linked glycosylation *in vitro*. We extended these studies *in vivo* by adoptively transferring leukemias into sublethally irradiated NSG mice which lack CD4^+^ and CD8α^+^ T lymphocytes but harbor mouse macrophages that are capable of phagocytosis (**Fig. 3D**) (*6*). Systemic administration of opsonizing anti-CD8α antibodies to sgCtrl-CD8α leukemia-bearing NSG mice *in vivo* resulted in a statistically significant prolongation in survival compared to mice that did not receive anti-CD8α antibodies (**Fig. S8I**). Notably, we observed that mice bearing sgC1GALT1-CD8α OE leukemias exhibited prolonged survival compared to NSG mice bearing sgCtrl-CD8α OE leukemias (**Fig. 3E**), demonstrating that the O-linked glycosylation pathway enables leukemia evasion of macrophages *in vivo*.

O-glycans often terminate in sialic acids, which are negatively charged sugar residues with immune inhibitory activity (*32*). As terminal sialic acids on cell surface glycoproteins are lost after genetic deletion of *C1GALT1* or *C1GALT1C1*, we enzymatically and genetically perturbed cell surface sialylation to determine if sialic acid modifications were responsible for inhibiting macrophage phagocytosis. First, we pre-treated both control and *C1GALT1*- or *C1GALT1C1*-deficient leukemias with *V. cholerae* sialidase, a pan-sialidase that removes all terminal sialic acid linkages (α2,3/α2,6/α2,8) (*33*). We hypothesized that enzymatic removal of terminal sialic acids from control leukemias would render them equally sensitive to phagocytosis as *C1GALT1-* or *C1GALT1C1*-deficient leukemias. Consistent with this hypothesis, increasing concentrations of sialidase abrogated the selective phagocytosis of *C1GALT1*- or *C1GALT1C1*-deficient leukemias in competition assays (**Fig. 3F**). Furthermore, selective treatment of control leukemia cells with a concentration of sialidase that completely removed cell surface sialic acids reversed the relative preference of human macrophages for *C1GALT1*- or *C1GALT1C1*-deficient leukemias that had not been treated with sialidase (**Fig. 3G**).

Sialic acid incorporation into glycans requires conversion of sialic acid to CMP-Sia by CMAS and the transport of CMP-Sia into the Golgi by SLC35A1 (*34*). Consistent with the sialidase result, knockout of *CMAS* or *SLC35A1* in leukemia cells enhanced phagocytosis (**Fig. S8J)**. These results demonstrate that the inhibitory effects of O-linked glycosylation on macrophage phagocytosis are mediated by the incorporation of terminal sialic acids to cell surface glycans.

### CD43 is the major O-linked glycoprotein that inhibits phagocytosis of leukemia

To determine the identity of the O-glycosylated cell surface proteins on leukemia cells that directly inhibit phagocytosis, we ranked all known cell surface and secreted proteins by mRNA expression and effect on macrophage phagocytosis. As O-glycosylation involves the addition of sugar molecules to a serine (Ser) or threonine (Thr), we also plotted the proportion of Ser/Thr residues for each protein, as each represents a potential O-glycosylation site (*35–37*). Strikingly, the mucin CD43 (*SPN*), whose extracellular domain is composed of >30% serine/threonine residues, scored the highest in this analysis (**Fig. 4A**).

**Fig. 4:**
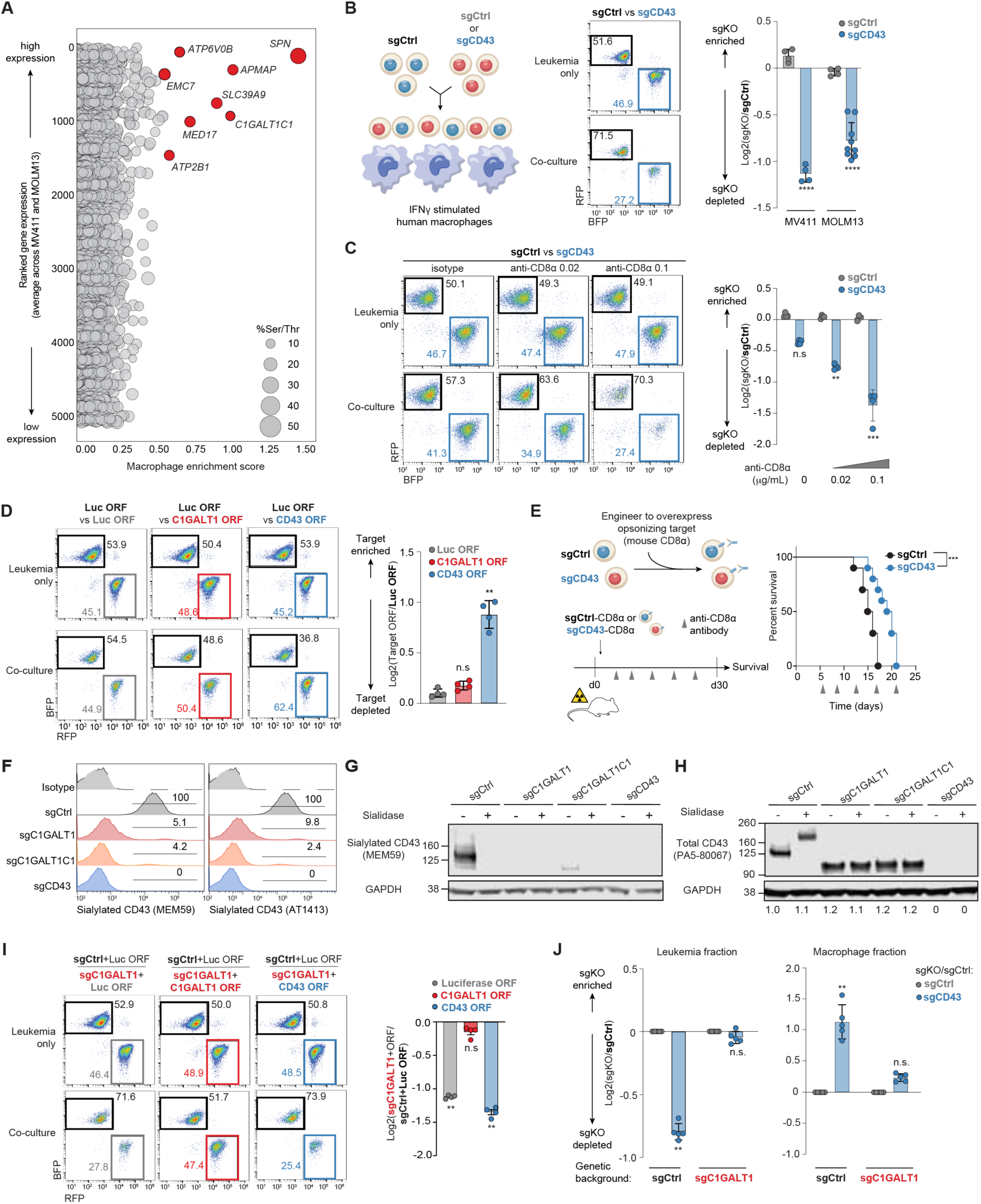
Sialylated CD43 is the major downstream effector of the O-linked glycosylation pathway and is sufficient to inhibit macrophage phagocytosis of human leukemia **a,** Scatter plot of functional impact in CRISPR AICP screens (*x*-axis) vs. relative gene expression (*y*-axis) of all known human cell surface proteins. Size of each dot is scaled to reflect the % of potential O-linked glycosylation sites. **b,** Schematic of competitive phagocytosis assay, representative flow cytometry plots, and bar graphs of relative phagocytosis of *SPN (CD43)-*deficient or control sgRNA versus control leukemia cells (MV411 and MOLM13) in coculture assays with IFNγ-stimulated macrophages. **c.** Representative flow cytometry plots and bar graphs of relative phagocytosis of *SPN (CD43)-*deficient or control leukemias overexpressing CD8α versus control MV411 leukemia cells overexpressing CD8α in coculture assays with varying doses of anti-CD8α opsonizing antibodies. **d.** Representative flow cytometry plots and bar graphs of relative phagocytosis of control leukemias overexpressing luciferase, C1GALT1, or CD43 versus control MV411 leukemia cells overexpressing luciferase in coculture assays with IFNγ-stimulated macrophages. **e**. Survival of mice challenged with *SPN (CD43)*-deficient or control MV411 leukemia cells overexpressing CD8α. Mice were intraperitoneally injected with systemic anti-CD8α antibodies starting on day +5. **f**. Representative flow plots for surface expression of sialylated CD43 in control, *C1GALT1*-deficient, *C1GALT1C1*-deficient, or *CD43*-deficient MV411 leukemias. **g.** Representative western blots for sialylated CD43 levels in MV411 leukemia cells treated without or with *V. cholerae* sialidase. **h.** Representative western blots for total CD43 levels in MV411 leukemia cells treated without or with *V. cholerae* sialidase **i.** Representative flow cytometry plots and bar graphs of relative phagocytosis of *C1GALT1*-deficient leukemias overexpressing luciferase, C1GALT1, or CD43 versus control MV411 leukemia cells overexpressing luciferase in coculture assays with IFNγ-stimulated macrophages. **j.** Relative phagocytosis of CD43-deficient or dual C1GALT1- and CD43-deficient leukemias after co-culture with IFNγ-stimulated macrophages. **b,c,d,h** Bar graphs indicate log (fold change) of the ratio of knockout cells relative to control after co-culture with IFNγ-stimulated macrophages. Data represent mean ± s.d. of four technical replicates and are representative of 3-5 independent experiments.

Genetic deletion of *SPN* (CD43) in MV411 and MOLM13 leukemias enhanced phagocytosis by IFNγ-stimulated macrophages in competition assays compared to control leukemias (**Fig. 4B**), while CD43-deficient leukemias engineered to overexpress CD8α (sgCD43-CD8α OE) were more sensitive to opsonizing antibody-mediated phagocytosis compared to sgCtrl-Luc leukemias **(Fig. 4C**). Conversely, CD43 overexpression in control MV411 leukemia cells, which have high endogenous expression of CD43, inhibited phagocytosis by IFNγ-stimulated macrophages *in vitro* in competition assays with control MV411 leukemias overexpressing the control antigen luciferase (**Fig. 4D**). In contrast, overexpression of *C1GALT1* in control leukemias had no impact on phagocytosis, suggesting that C1GALT1 enzymatic activity is not rate-limiting in leukemias (**Fig. 4D**). Furthermore, genetic deletion of CD43 in CD8α OE leukemias resulted in a statistically significant prolongation of survival *in vivo* in mice treated with anti-CD8α compared to sgCtrl-CD8α OE leukemias (**Fig. 4E**). Similar to our observations with *C1GALT1*- or *C1GALT1C1*-deficient leukemias, CD43-deficient leukemias exhibited no growth defects *in vitro* (**Fig. S9A**). Thus, CD43 expression on leukemia cells inhibits macrophage phagocytosis both *in vitro* and i*n vivo*.

We hypothesized that loss of O-glycosylation impairs sialylation of CD43 and measured sialylated CD43 using an anti-CD43 antibody (clone MEM59) with known specificity for sialylated but not desialylated CD43 (*38*, *39*). As expected, we did not detect sialylated CD43 on CD43-deficient leukemias or control leukemia cells treated with sialidase **(Fig. S9B**). We observed near-complete loss of sialylated CD43 in C1GALT1- or C1GALT1C1-deficient leukemias, both by flow cytometry and western blot (**Fig. 4F, left; Fig. 4G**). We verified that loss of O-linked glycosylation impairs CD43 sialylation with a second antibody specific for sialylated but not desialylated human CD43 (clone AT1413) **(Fig. 4F, right)** (*40*, *41*). In contrast, we saw no changes in total CD43 levels in *C1GALT1*- or *C1GALT1C1*-deficient leukemias, but observed a mobility shift consistent with loss of O-glycans (**Fig. 4H**) (*42*, *43*). While overexpression of C1GALT1 in C1GALT1-deficient leukemias completely reversed their susceptibility to macrophage phagocytosis, overexpression of CD43 in C1GALT1-deficient leukemias had no effect (**Fig. 4I**), consistent with the hypothesis that intact sialylation of CD43 is required for inhibition of phagocytosis and this deficiency cannot be rescued simply by increasing the total expression level of CD43. Thus, O-linked glycosylation inhibits macrophage phagocytosis of leukemia through post-translational addition of inhibitory terminal sialic acid residues to CD43.

To determine whether CD43 is the primary mediator of the inhibitory effect of O-glycosylation on macrophage phagocytosis, we performed genetic epistasis experiments by deleting C1GALT1, CD43, or both genes in leukemia cells prior to co-culture with macrophages. As expected, genetic loss of CD43 in a control background (*sgCtrl*) strongly enhanced phagocytosis (**Fig. 4J**). However, in cells that lacked C1GALT1 or its chaperone C1GALT1C1 (*sgC1GALT1, sgC1GALT1C1*), genetic loss of CD43 had no effect, suggesting that the inhibitory effect of CD43 is absolutely dependent on an intact O-glycosylation pathway (**Fig. 4J**, **Fig. S9C**). Thus, sialylated CD43 is the major downstream effector through which C1GALT1 and C1GALT1C1 inhibit macrophage phagocytosis.

### CD43 inhibition of phagocytosis is not mediated by the known sialic acid-binding lectins SIGLEC-7 or SIGLEC-9

SIGLECs are a family of inhibitory sialic acid-binding cell surface receptors that are expressed by several mature immune cells, including monocytes, granulocytes, macrophages, and natural killer (NK) cells (*44*– *46*). We hypothesized that SIGLEC receptors on human macrophages deliver inhibitory signals from sialylated CD43 to restrain phagocytosis. We detected abundant expression of several SIGLEC family members in M-CSF differentiated human macrophages, including SIGLEC-7 and SIGLEC-9 (**Fig. S10A-B, Table S11**). As SIGLEC-7 and SIGLEC-9 on macrophages have been implicated as functionally important sensors of sialylation (*47–49*), we assessed whether sialylated CD43 on leukemia cells inhibits phagocytosis through SIGLEC-7 and/or SIGLEC-9. Surprisingly, genetic loss of CD43 in leukemias did not impact binding of either SIGLEC as measured by recombinant SIGLEC-7 or SIGLEC-9 Fc staining (**Fig. S10c)**. Furthermore, neither genetic deletion of both SIGLEC-7 and SIGLEC-9 in macrophages nor dual antibody blockade with anti-SIGLEC-7/anti-SIGLEC-9 neutralizing antibodies impacted phagocytosis (**Fig. S10D-F**). Thus, SIGLEC-7 and SIGLEC-9 are not necessary for mediating the inhibitory effects of sialylated CD43 on human macrophage phagocytosis. In addition to binding to SIGLEC receptors, sialylated O-glycans can impact biophysical interactions through steric or electrostatic hindrance. CD43 has a heavily sialylated rod-like ectodomain that extends outwards from the plasma membrane and imparts a negative charge (*50*). To assess whether the biophysical properties of CD43 can impair intercellular interactions, we performed cell:cell interaction avidity measurements using macrophages co-cultured with either control or CD43-deficient leukemias. CD43-deficient leukemias bound human macrophages with higher avidity than control leukemias (**Fig. S10G).** These results suggest that sialylated CD43 may restrain macrophage phagocytosis through modulation of macrophage:leukemia binding. Further investigation is required to identify whether this is through a SIGLEC-dependent or SIGLEC-independent mechanism.

### Anti-CD43 antibodies as a therapeutic strategy for AML

To explore the suitability of CD43 as a therapeutic target in AML, we examined its expression in patient-derived normal and malignant hematopoietic cells. We characterized the expression of CD43 in primary AML patient samples using single-cell RNA sequencing (scRNA-seq) to avoid the confounding effects of mixtures of malignant and normal myeloid cells. First, we re-analyzed a scRNA-seq dataset of bone marrow aspirates from healthy donors and AML patients and compared the expression of CD43 (*SPN)*(*9*). *SPN* expression was highest in malignant hematopoietic stem cell (HSC)-like and malignant hematopoietic progenitor cell (HPC)-like cells compared to normal HSC-like and HPC-like cells (**Fig. 5A-C**). Western blot analysis also showed that expression of sialylated CD43 was higher in AML compared to normal peripheral blood mononuclear cells or purified monocytes, and we confirmed high surface expression in AML patients by flow cytometry (**Fig. 5D-E**). In contrast, expression of O-glycosylation enzymes (*C1GALT1, C1GALT1C1, SLC39A9, SLC35A2)* was similar between normal and malignant hematopoietic cells (**Fig. S11A**). These results indicate that CD43 is overexpressed in human leukemia cells relative to normal hematopoietic cells, making it attractive as a therapeutic target in AML.

**Fig. 5:**
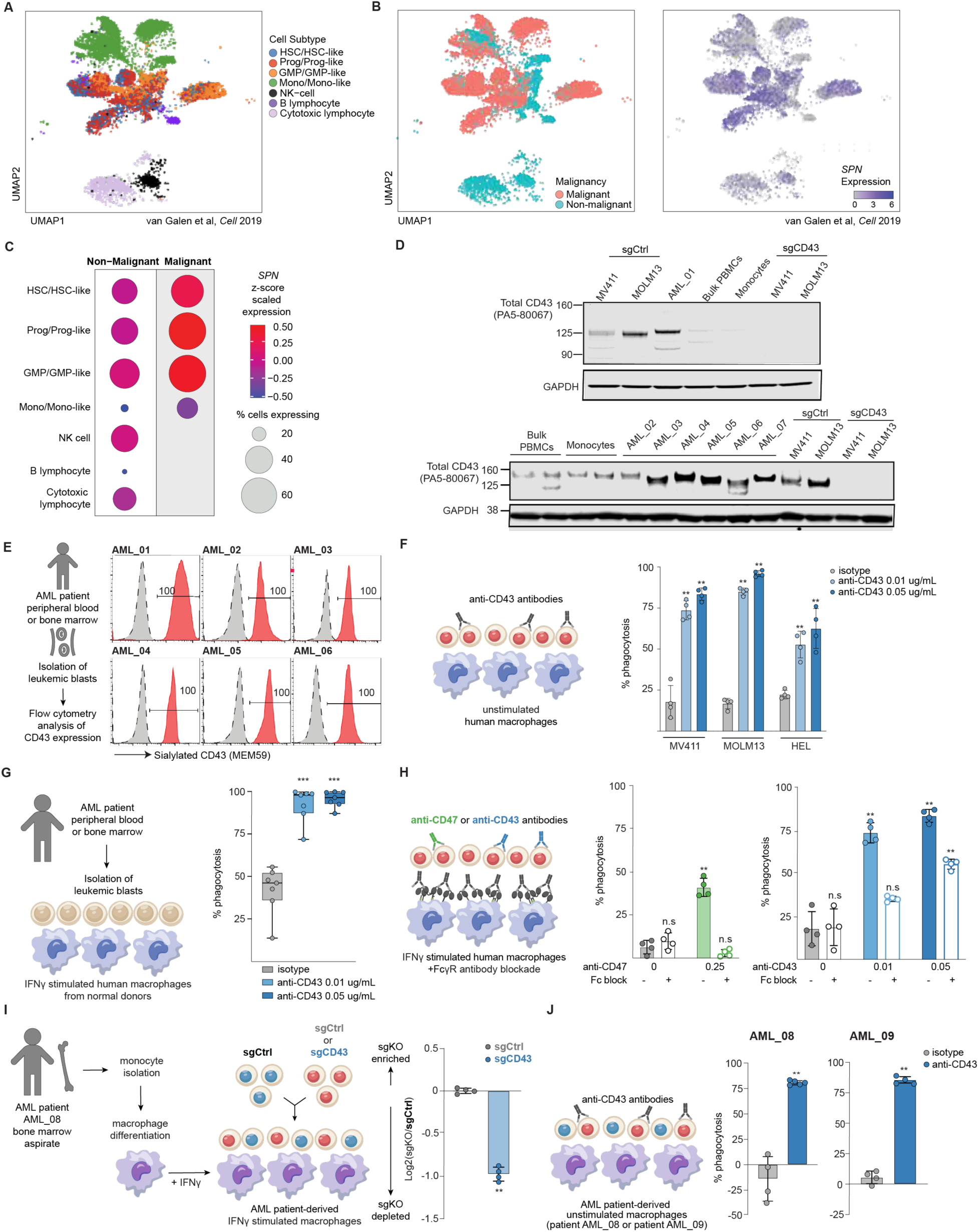
anti-CD43 antibodies enhance human macrophage phagocytosis of leukemia cell lines and primary patient samples. **a-b,** Uniform manifold approximation and projection (UMAP) of single cell RNA-sequencing (scRNA-seq) profiles of bone marrow aspirate cells taken from healthy patients or AML patients. Expression of SPN/CD43 is shown on the far right panel. **c,** Dot plot summary of z-score scaled expression of CD43 across normal and malignant cell subtypes. Size of each dot is scaled to reflect the absolute % of cells in which SPN transcripts were detected. **d,** Representative western blots for total CD43 expression in human leukemia cell lines (MV411, MOLM13), primary AML blasts (AML_01 - AML07), and normal immune cell subsets (bulk PBMCs, monocytes). **e,** Flow cytometry plots for sialylated CD43 expression on fresh blasts from AML patients (AML01 - AML06) **f,** Schematic of phagocytosis experiments with anti-CD43 antibodies and bar graphs of absolute phagocytosis of human leukemia cell lines (MV411, MOLM13, HEL) after addition of various doses of anti-CD43 antibodies to macrophage co-culture assays **g,** Schematic of phagocytosis assays of patient-derived AML blasts and bar graphs of absolute phagocytosis of primary AML cells after addition of varying doses of anti-CD43 antibodies to macrophage co-culture assays **h,** Schematic of phagocytosis assays with FcR blockade prior to addition of either anti-CD47 or anti-CD43 antibodies and bar graphs of absolute phagocytosis of MV411 leukemias with or without Fc blockade prior to addition of anti-CD47 or anti-CD43 antibodies. **i,** Schematic of phagocytosis assays with AML patient-derived macrophages and bar graphs of relative phagocytosis of control or CD43-deficient leukemias versus control MV411 leukemias **j,** Schematic of phagocytosis assays with AML patient-derived macrophages and bar plots of absolute phagocytosis of MV411 leukemias with or without anti-CD43 antibodies. **f,g,h,i,j** Data represent mean ± s.d. of four technical replicates and are representative of 3-5 independent experiments.

Given the high expression of sialylated CD43 in both human leukemia cells and primary AML blasts, we evaluated the impact of targeting sialylated CD43 with antibodies. We observed that the addition of anti-CD43 antibodies with high specificity for sialylated CD43 (clones MEM59 and 10G7) to co-culture assays enhanced phagocytosis of MV411, MOLM13, and HEL AML cell lines in a dose-dependent manner, as measured by flow cytometry or fluorescence microscopy (**Fig. 5F Fig. S11B-C**). We extended these experiments beyond AML cell lines to primary patient-derived AML blasts, and similarly observed that anti-CD43 antibody treatment enhanced their phagocytosis (**Fig. 5G**). Taken together, these results indicate that CD43 antibody blockade phenocopies genetic loss of CD43.

Anti-CD43 antibodies might enhance macrophage phagocytosis through at least two distinct mechanisms: 1) direct interference with the inhibitory effects of sialylated CD43 on human macrophages; 2) activation of the FcγR after opsonization of leukemia cells with anti-CD43 antibodies. If anti-CD43 antibodies act entirely through activation of the FcγR, then abrogating the ability of anti-CD43 antibodies to bind the FcγR should reverse the enhancement of phagocytosis after CD43 blockade. We observed that anti-CD43 antibodies with impaired binding to FcγR (D265A mouse IgG1 “Fc dead”)(*51*) still enhanced phagocytosis in a dose-dependent manner (**Fig. S11D**). Furthermore, unlike anti-CD47 antibodies, anti-CD43 antibodies enhanced macrophage phagocytosis even after blockade of all FcγR (CD16/CD32/CD64) on human macrophages with neutralizing antibodies (**Fig. 5H**). Therefore, unlike anti-CD47 antibodies, anti-CD43 antibodies enhance human macrophage phagocytosis through a combination of FcγR-independent and FcγR-dependent mechanisms. Because myeloid cell dysfunction has been observed in patients with AML (*9*), we next tested whether macrophages derived from patients with active AML were responsive to anti-CD43 mediated enhanced phagocytosis. AML patient-derived macrophages showed increased phagocytosis either following CD43 antibody blockade or knockout of CD43 in MV411 cells (**Fig. 5I-J**). These results indicate that AML patient-derived macrophages retain their ability to respond to loss of sialylated CD43, resulting in increased phagocytosis.

## Discussion

Macrophages are a critical effector population of innate immunity and are highly abundant in the tumor microenvironment in most types of cancer (*52–54*). In the context of certain activation cues, they can exert potent anti-tumor activity through cytokine production, antigen presentation to T cells, and importantly, direct phagocytosis and elimination of tumor cells. The importance of this effector function for successful anti-tumor immune responses is now broadly appreciated and has led to the development of macrophage targeting drugs and cell therapies (*25*, *55*). Here, we used genome-scale CRISPR screens in co-culture assays of human leukemias and human monocyte-derived macrophages to discover the pathways that restrain human macrophage antibody-independent and antibody-dependent phagocytosis. Our screens recovered previously identified regulators of phagocytosis such as APMAP (*11*), confirming the reliability of the screening approach. Surprisingly, this study revealed that genetic loss of CD47 alone is insufficient to enhance phagocytosis by human macrophages, whereas genetic loss of CD47 does stimulate phagocytosis by murine macrophages, as has been previously reported (*2*, *11*). Additionally, a *Sirpa* polymorphism in NSG mice that confers strong recognition of human CD47 (*56*) may in part explain encouraging results in pre-clinical leukemia xenograft models (*2*, *3*) that contrast with the lack of clinical benefit of CD47 blocking antibodies in recent phase 3 trials (*4*)(*2*, *3*). We also note that our discovery of the MHC-I:LILRB1/2 axis as the dominant regulator of antibody-dependent cellular phagocytosis was missed in prior studies due to species-mismatched co-culture assays (*11*). It remains possible that CD47-blocking antibodies could have therapeutic benefit through ADCP, as opposed to only serving as a macrophage-inhibitory checkpoint as originally proposed. More generally, our results underscore the importance of testing therapeutic hypotheses in human experimental systems whenever possible.

Most importantly, our study revealed that O-linked glycosylation and sialylation restrain both antibody-independent and antibody-dependent phagocytosis of leukemia, largely through sialylation of the surface glycoprotein CD43. Genetic deletion or antibody targeting sialylated CD43 enhanced phagocytosis of primary AML blasts by macrophages from healthy donors and AML patients.

O-glycan modifications that terminate in sialic acid, particularly on the highly expressed cell surface mucin CD43, potently inhibited macrophage phagocytosis. Overexpression of CD43 in leukemia cells that lacked the ability to undergo mucin-type O-linked glycosylation had no impact on macrophage phagocytosis, suggesting that post-translational glyco-modifications (and not the underlying protein structure) are responsible for mediating immune evasion. SIGLEC receptors are an important class of sialic acid-binding inhibitory immune receptors (*44*), and previous work has implicated SIGLEC-7 as a functionally important cognate receptor for sialylated CD43 (*46*). However, in our studies neither genetic deletion of SIGLEC-7, or the related receptor SIGLEC-9, in human macrophages nor antibody-mediated blockade of SIGLEC-7/9 abrogated the enhancement of phagocytosis of C1GALT1-deficient or CD43-deficient leukemias. Similar observations were recently reported with potent SIGLEC-7/SIGLEC-9 dual degraders, which exhibited no impact on human macrophage phagocytosis (*57*). This suggests that loss of CD43 may enhance phagocytosis either through other inhibitory receptors with carbohydrate recognition domains (galectins, C-type lectins, or other SIGLECs) or perhaps through a receptor-independent mechanism. CD43 has >30 reported extracellular O-linked glycosylation sites, and extends 45 nm from the cell surface (*50*). This large, negatively charged structure could form a barrier that sterically or electrostatically hinders high avidity interactions between leukemia cells and immune effector cells (*58*). Further studies will be required to understand the mechanism by which sialylated CD43 restricts phagocytosis by macrophages.

In conclusion, our study highlights the importance of sialylated CD43 as a regulator of macrophage phagocytosis of AML, and suggests that the development of CD43-directed therapies should be explored further. While CD43 is predominantly expressed in hematopoietic malignancies, there may be other heavily sialylated mucin and mucin-like cell surface proteins that play a similar role in restricting the phagocytic activity of nearby macrophages in solid tumors (*59*). For example, antibody conjugated sialidases targeting solid tumors have demonstrated enhanced anti-tumor immunity highlighting a role for identifying and targeting sialoglycans in a diverse range of tumor types (*60*, *61*). Thus, O-linked glycosylation and sialylation of the cell surface represents a glyco-immune checkpoint that restrains phagocytosis by macrophages and likely has immune inhibitory activity that extends to a broad range of tumor contexts and immune interactions.

## Acknowledgments

We thank all members of the Manguso and Golub labs for discussions and feedback. We thank Abigail Marola, Amyah Harris, and the rest of the MGH Tissue Bank team for providing AML patient samples. We thank P. Rogers and Broad Technology Space for assistance with live cell imaging. We also thank T. Caron and the Broad vivarium staff for their support and assistance. Research reported in this publication was supported by the Ted and Eileen Pasquarello Tissue Bank in Hematologic Malignancies.

## Finding

National Institutes of Health grant R35 CA242457 (TRG) Calico Life Sciences LLC. (RTM, TRG)

National Institutes of Health grant T32 CA009172-46 (JC) National Institutes of Health grant T32 HL116324-07 (MV)

Damon Runyon Cancer Research Foundation David M. Livingston, MD Physician-Scientist (MV) Edward P. Evans Foundation EvansMDS Young Investigator (MV)

Polish National Agency for Academic Exchange – the Walczak Programme (EW)

## Author contributions

Conceptualization: J.C., M.V., T.R.G., R.T.M

Formal analysis: J.C., M.V., Y.L., C.N., J.P., D.W., A.C., A.K., M.K.

Funding acquisition: K.B.Y., T.R.G., R.T.M

Investigation: J.C., M.V., S.N., G.S., E.W., M.H., S.L., M.S., I.S., E.C.W., L.C., F.B., O.I.A.

Methodology: J.C., M.V., K.B.Y., R.T.M

Project administration: K.B.Y., T.R.G., R.T.M

Resources: J.G.D., D.P., L.T., E.W., D.D., J.G., R.M.S., T.A.G Supervision: J.C., M.V., K.B.Y., T.R.G., R.T.M

Visualization: J.C., M.V., S.N., Y.L., C.N., J.P, A.C

Writing - original draft: J.C., M.V., K.B.Y., R.T.M

Writing - review and editing: J.C., M.V., K.B.Y., T.R.G., R.T.M

## Data and materials availability

All data are available in the manuscript or the supplementary materials. Further information and requests for resources and reagents should be directed to and will be fulfilled by the corresponding authors. Cell lines and DNA constructs generated in this study may be available upon request. Human macrophage RNA-seq data has been deposited in Gene Expression Omnibus under accession code GSEXXXXXX.

## Supplementary Materials

Materials and Methods

Supplementary Text Figs. S1 to S11

Tables S1 to S11

## References

1. G. C. Brown, Cell death by phagocytosis. Nat. Rev. Immunol. 24, 91–102 (2024).

2. S. Jaiswal, C. H. M. Jamieson, W. W. Pang, C. Y. Park, M. P. Chao, R. Majeti, D. Traver, N. van Rooijen, I. L. Weissman, CD47 is upregulated on circulating hematopoietic stem cells and leukemia cells to avoid phagocytosis. Cell 138, 271–285 (2009).

3. R. Majeti, M. P. Chao, A. A. Alizadeh, W. W. Pang, S. Jaiswal, K. D. Gibbs Jr, N. van Rooijen, I. L. Weissman, CD47 is an adverse prognostic factor and therapeutic antibody target on human acute myeloid leukemia stem cells. Cell 138, 286–299 (2009).

4. ENHANCE, 2, 3 studies document for posting 26 Feb. https://www.gilead.com/-/media/files/pdfs/other/magrolimab-trials-summary.pdf.

5. J. Mestas, C. C. W. Hughes, Of mice and not men: differences between mouse and human immunology. J. Immunol. 172, 2731–2738 (2004).

6. K. Takenaka, T. K. Prasolava, J. C. Y. Wang, S. M. Mortin-Toth, S. Khalouei, O. I. Gan, J. E. Dick, J. S. Danska, Polymorphism in Sirpa modulates engraftment of human hematopoietic stem cells. Nat. Immunol. 8, 1313–1323 (2007).

7. N. Daver, G. Garcia-Manero, S. Basu, P. C. Boddu, M. Alfayez, J. E. Cortes, M. Konopleva, F. Ravandi-Kashani, E. Jabbour, T. Kadia, G. M. Nogueras-Gonzalez, J. Ning, N. Pemmaraju, C. D. DiNardo, M. Andreeff, S. A. Pierce, T. Gordon, S. M. Kornblau, W. Flores, Z. Alhamal, C. Bueso-Ramos, J. L. Jorgensen, K. P. Patel, J. Blando, J. P. Allison, P. Sharma, H. Kantarjian, Efficacy, safety, and biomarkers of response to azacitidine and nivolumab in relapsed/refractory acute myeloid leukemia: A nonrandomized, open-label, phase II study. Cancer Discov. 9, 370–383 (2019).

8. J. P. Bewersdorf, R. M. Shallis, E. Sharon, S. Park, R. Ramaswamy, C. E. Roe, J. M. Irish, A. Caldwell, W. Wei, A. Yacoub, Y. F. Madanat, J. F. Zeidner, J. K. Altman, O. Odenike, S. Yerrabothala, T. Kovacsovics, N. A. Podoltsev, S. Halene, R. F. Little, R. Piekarz, S. D. Gore, T. K. Kim, A. M. Zeidan, A multicenter phase Ib trial of the histone deacetylase inhibitor entinostat in combination with pembrolizumab in patients with myelodysplastic syndromes/neoplasms or acute myeloid leukemia refractory to hypomethylating agents. Ann. Hematol. 103, 105–116 (2024).

9. P. van Galen, V. Hovestadt, M. H. Wadsworth Ii, T. K. Hughes, G. K. Griffin, S. Battaglia, J. A. Verga, J. Stephansky, T. J. Pastika, J. Lombardi Story, G. S. Pinkus, O. Pozdnyakova, I. Galinsky, R. M. Stone, T. A. Graubert, A. K. Shalek, J. C. Aster, A. A. Lane, B. E. Bernstein, Single-cell RNA-seq reveals AML hierarchies relevant to disease progression and immunity. Cell 176, 1265–1281.e24 (2019).

10. X. Huang, Y. Li, M. Fu, H.-B. Xin, Polarizing macrophages in vitro. Methods Mol. Biol. 1784, 119–126 (2018).

11. R. A. Kamber, Y. Nishiga, B. Morton, A. M. Banuelos, A. A. Barkal, F. Vences-Catalán, M. Gu, D. Fernandez, J. A. Seoane, D. Yao, K. Liu, S. Lin, K. Spees, C. Curtis, L. Jerby-Arnon, I. L. Weissman, J. Sage, M. C. Bassik, Inter-cellular CRISPR screens reveal regulators of cancer cell phagocytosis. Nature 597, 549–554 (2021).

12. M. E. W. Logtenberg, J. H. M. Jansen, M. Raaben, M. Toebes, K. Franke, A. M. Brandsma, H. L. Matlung, A. Fauster, R. Gomez-Eerland, N. A. M. Bakker, S. van der Schot, K. A. Marijt, M. Verdoes, J. B. A. G. Haanen, J. H. van den Berg, J. Neefjes, T. K. van den Berg, T. R. Brummelkamp, J. H. W. Leusen, F. A. Scheeren, T. N. Schumacher, Glutaminyl cyclase is an enzymatic modifier of the CD47-SIRPα axis and a target for cancer immunotherapy. Nat. Med. 25, 612–619 (2019).

13. J. C. Osorio, P. Smith, D. A. Knorr, J. V. Ravetch, The antitumor activities of anti-CD47 antibodies require Fc-FcγR interactions. Cancer Cell 41, 2051–2065.e6 (2023).

14. S. Jain, A. Van Scoyk, E. A. Morgan, A. Matthews, K. Stevenson, G. Newton, F. Powers, A. Autio, A. Louissaint, G. Pontini, J. C. Aster, F. W. Luscinskas, D. M. Weinstock, Targeted inhibition of CD47-SIRPα requires Fc-FcγR interactions to maximize activity in T-cell lymphomas. Blood 134, 1430–1440 (2019).

15. S. T. Jung, W. Kelton, T. H. Kang, D. T. W. Ng, J. T. Andersen, I. Sandlie, C. A. Sarkar, G. Georgiou, Effective phagocytosis of low Her2 tumor cell lines with engineered, aglycosylated IgG displaying high FcγRIIa affinity and selectivity. ACS Chem. Biol. 8, 368–375 (2013).

16. L.-C. Tsao, E. J. Crosby, T. N. Trotter, J. Wei, T. Wang, X. Yang, A. N. Summers, G. Lei, C. A. Rabiola, L. A. Chodosh, W. J. Muller, H. K. Lyerly, Z. C. Hartman, Trastuzumab/pertuzumab combination therapy stimulates antitumor responses through complement-dependent cytotoxicity and phagocytosis. JCI Insight 7 (2022).

17. M. Colonna, H. Nakajima, F. Navarro, M. López-Botet, A novel family of Ig-like receptors for HLA class I molecules that modulate function of lymphoid and myeloid cells. J. Leukoc. Biol. 66, 375–381 (1999).

18. M. Shiroishi, K. Tsumoto, K. Amano, Y. Shirakihara, M. Colonna, V. M. Braud, D. S. J. Allan, A. Makadzange, S. Rowland-Jones, B. Willcox, E. Y. Jones, P. A. van der Merwe, I. Kumagai, K. Maenaka, Human inhibitory receptors Ig-like transcript 2 (ILT2) and ILT4 compete with CD8 for MHC class I binding and bind preferentially to HLA-G. Proc. Natl. Acad. Sci. U. S. A. 100, 8856–8861 (2003).

19. M. Colonna, F. Navarro, T. Bellón, M. Llano, P. García, J. Samaridis, L. Angman, M. Cella, M. López-Botet, A common inhibitory receptor for major histocompatibility complex class I molecules on human lymphoid and myelomonocytic cells. J. Exp. Med. 186, 1809–1818 (1997).

20. M. Colonna, J. Samaridis, M. Cella, L. Angman, R. L. Allen, C. A. O’Callaghan, R. Dunbar, G. S. Ogg, V. Cerundolo, A. Rolink, Human myelomonocytic cells express an inhibitory receptor for classical and nonclassical MHC class I molecules. J. Immunol. 160, 3096–3100 (1998).

21. Regulation of immune disorders by PIR-B, a critical inhibitory Ig-like receptor for MHC class I molecules.

22. T. Takai, A novel recognition system for MHC class I molecules constituted by PIR. Adv. Immunol. 88, 161–192 (2005).

23. A. Barkal, K. Weiskopf, K. S. Kao, S. R. Gordon, B. Rosental, Y. Y. Yiu, B. M. George, M. Markovic, N. G. Ring, J. M. Tsai, K. M. McKenna, P. Y. Ho, R. Z. Cheng, J. Y. Chen, L. J. Barkal, A. M. Ring, I. L. Weissman, R. L. Maute, Engagement of MHC class I by the inhibitory receptor LILRB1 suppresses macrophages and is a target of cancer immunotherapy. Nat. Immunol. 19, 76–84 (2018).

24. J. Tian, A. M. Ashique, S. Weeks, T. Lan, H. Yang, H.-I. H. Chen, C. Song, K. Koyano, K. Mondal, D. Tsai, I. Cheung, M. Moshrefi, A. Kekatpure, B. Fan, B. Li, S. Qurashi, L. Rocha, J. Aguayo, C. Rodgers, M. Meza, D. Heeke, S. M. Medfisch, C. Chu, S. Starck, N. P. Basak, S. Sankaran, M. Malhotra, S. Crawley, T.-T. Tran, D. Y. Duey, C. Ho, I. Mikaelian, W. Liu, L. B. Rivera, J. Huang, K. J. Paavola, K. O’Hollaren, L. K. Blum, V. Y. Lin, P. Chen, A. Iyer, S. He, J. M. Roda, Y. Wang, J. Sissons, A. K. Kutach, D. D. Kaplan, G. W. Stone, ILT2 and ILT4 drive myeloid suppression via both overlapping and distinct mechanisms. Cancer Immunol. Res. 12, 592–613 (2024).

25. A Study of OR502, a Monoclonal Antibody Targeting LILRB2, Alone and in Combination With Anticancer Agents, NCT06090266. https://clinicaltrials.gov/study/NCT06090266?intr=LILRB2&rank=1.

26. R. W. Lentz, M. D. Colton, S. S. Mitra, W. A. Messersmith, Innate immune checkpoint inhibitors: The next breakthrough in medical oncology? Mol. Cancer Ther. 20, 961–974 (2021).

27. Y. Zhou, M. Fei, G. Zhang, W.-C. Liang, W. Lin, Y. Wu, R. Piskol, J. Ridgway, E. McNamara, H. Huang, J. Zhang, J. Oh, J. M. Patel, D. Jakubiak, J. Lau, B. Blackwood, D. D. Bravo, Y. Shi, J. Wang, H.-M. Hu, W. P. Lee, R. Jesudason, D. Sangaraju, Z. Modrusan, K. R. Anderson, S. Warming, M. Roose-Girma, M. Yan, Blockade of the phagocytic receptor MerTK on tumor-associated macrophages enhances P2X7R-dependent STING activation by tumor-derived cGAMP. Immunity 52, 357–373.e9 (2020).

28. V. A. Fadok, D. R. Voelker, P. A. Campbell, J. J. Cohen, D. L. Bratton, P. M. Henson, Exposure of phosphatidylserine on the surface of apoptotic lymphocytes triggers specific recognition and removal by macrophages. J. Immunol. 148, 2207–2216 (1992).

29. R. S. Scott, E. J. McMahon, S. M. Pop, E. A. Reap, R. Caricchio, P. L. Cohen, H. S. Earp, G. K. Matsushima, Phagocytosis and clearance of apoptotic cells is mediated by MER. Nature 411, 207–211 (2001).

30. Y. Wang, T. Ju, X. Ding, B. Xia, W. Wang, L. Xia, M. He, R. D. Cummings, Cosmc is an essential chaperone for correct protein O-glycosylation. Proc. Natl. Acad. Sci. U. S. A. 107, 9228–9233 (2010).

31. T. B. Rømer, F. Khoder-Agha, M. K. M. Aasted, N. de Haan, S. Horn, A. Dylander, T. Zhang, E. M. H. Pallesen, S. Dabelsteen, M. Wuhrer, C. F. Høgsbro, E. A. Thomsen, J. G. Mikkelsen, H. H. Wandall, CRISPR-screen identifies ZIP9 and dysregulated Zn2+ homeostasis as a cause of cancer-associated changes in glycosylation. Glycobiology 33, 700–714 (2023).

32. Varki, P. Gagneux, Multifarious roles of sialic acids in immunity: Roles of sialic acids in immunity. Ann. N. Y. Acad. Sci. 1253, 16–36 (2012).

33. P. Corfield, H. Higa, J. C. Paulson, R. Schauer, The specificity of viral and bacterial sialidases for alpha(2-3)- and alpha(2-6)-linked sialic acids in glycoproteins. Biochim. Biophys. Acta 744, 121–126 (1983).

34. T. K. Altheide, T. Hayakawa, T. S. Mikkelsen, S. Diaz, N. Varki, A. Varki, System-wide genomic and biochemical comparisons of sialic acid biology among primates and rodents: Evidence for two modes of rapid evolution. J. Biol. Chem. 281, 25689–25702 (2006).

35. S. L. King, H. J. Joshi, K. T. Schjoldager, A. Halim, T. D. Madsen, M. H. Dziegiel, A. Woetmann, S. Y. Vakhrushev, H. H. Wandall, Characterizing the O-glycosylation landscape of human plasma, platelets, and endothelial cells. Blood Adv. 1, 429–442 (2017).

36. W. Yang, M. Ao, Y. Hu, Q. K. Li, H. Zhang, Mapping the O-glycoproteome using site-specific extraction of O-linked glycopeptides (EXoO). Mol. Syst. Biol. 14, e8486 (2018).

37. A. Yamada, M. Shiota, D. Shinmachi, T. Ono, S. Tsuchiya, M. Hosoda, A. Fujita, N. P. Aoki, Y. Watanabe, N. Fujita, K. Angata, H. Kaji, H. Narimatsu, S. Okuda, K. F. Aoki-Kinoshita, The GlyCosmos Portal: a unified and comprehensive web resource for the glycosciences. Nat. Methods 17, 649–650 (2020).

38. W. de Smet, H. Walter, L. van Hove, A new CD43 monoclonal antibody induces homotypic aggregation of human leucocytes through a CD11a/CD18-dependent and -independent mechanism. Immunology 79, 46–54 (1993).

39. M. Alvarado, C. Klassen, J. Cerny, V. Horejsí, R. E. Schmidt, MEM-59 monoclonal antibody detects a CD43 epitope involved in lymphocyte activation. Eur. J. Immunol. 25, 1051–1055 (1995).

40. M. A. Gillissen, G. de Jong, M. Kedde, E. Yasuda, S. E. Levie, G. Moiset, P. J. Hensbergen, A. Q. Bakker, K. Wagner, J. Villaudy, P. M. van Helden, H. Spits, M. D. Hazenberg, Patient-derived antibody recognizes a unique CD43 epitope expressed on all AML and has antileukemia activity in mice. Blood Adv. 1, 1551–1564 (2017).

41. L. Bartels, G. de Jong, M. A. Gillissen, E. Yasuda, V. Kattler, C. Bru, C. Fatmawati, S. E. van Hal-van Veen, M. G. Cercel, G. Moiset, A. Q. Bakker, P. M. van Helden, J. Villaudy, M. D. Hazenberg, H. Spits, K. Wagner, A chemo-enzymatically linked bispecific antibody retargets T cells to a sialylated Epitope on CD43 in acute myeloid leukemia. Cancer Res. 79, 3372–3382 (2019).

42. S. R. Carlsson, M. Fukuda, Isolation and characterization of leukosialin, a major sialoglycoprotein on human leukocytes. J. Biol. Chem. 261, 12779–12786 (1986).

43. O. Saitoh, F. Piller, R. I. Fox, M. Fukuda, T-lymphocytic leukemia expresses complex, branched O-linked oligosaccharides on a major sialoglycoprotein, leukosialin. Blood 77, 1491–1499 (1991).

44. P. R. Crocker, J. C. Paulson, A. Varki, Siglecs and their roles in the immune system. Nat. Rev. Immunol. 7, 255–266 (2007).

45. R. M. Wen, J. C. Stark, G. E. W. Marti, Z. Fan, A. Lyu, F. J. Garcia Marques, X. Zhang, N. M. Riley, S. M. Totten, A. Bermudez, R. Nolley, H. Zhao, L. Fong, E. G. Engleman, S. J. Pitteri, C. R. Bertozzi, J. D. Brooks, Sialylated glycoproteins suppress immune cell killing by binding to Siglec-7 and Siglec-9 in prostate cancer. J. Clin. Invest. 134 (2024).

46. S. Wisnovsky, L. Möckl, S. A. Malaker, K. Pedram, G. T. Hess, N. M. Riley, M. A. Gray, B. A. H. Smith, M. C. Bassik, W. E. Moerner, C. R. Bertozzi, Genome-wide CRISPR screens reveal a specific ligand for the glycan-binding immune checkpoint receptor Siglec-7. Proc. Natl. Acad. Sci. U. S. A. 118, e2015024118 (2021).

47. A. Ibarlucea-Benitez, P. Weitzenfeld, P. Smith, J. V. Ravetch, Siglecs-7/9 function as inhibitory immune checkpoints in vivo and can be targeted to enhance therapeutic antitumor immunity. Proc. Natl. Acad. Sci. U. S. A. 118, e2107424118 (2021).

48. E. Rodriguez, K. Boelaars, K. Brown, R. J. Eveline Li, L. Kruijssen, S. C. M. Bruijns, T. van Ee, S. T. T. Schetters, M. H. W. Crommentuijn, J. C. van der Horst, N. C. T. van Grieken, S. J. van Vliet, G. Kazemier, E. Giovannetti, J. J. Garcia-Vallejo, Y. van Kooyk, Sialic acids in pancreatic cancer cells drive tumour-associated macrophage differentiation via the Siglec receptors Siglec-7 and Siglec-9. Nat. Commun. 12, 1270 (2021).

49. P. Schmassmann, J. Roux, A. Buck, N. Tatari, S. Hogan, J. Wang, N. Rodrigues Mantuano, R. Wieboldt, S. Lee, B. Snijder, D. Kaymak, T. A. Martins, M.-F. Ritz, T. Shekarian, M. McDaid, M. Weller, T. Weiss, H. Läubli, G. Hutter, Targeting the Siglec-sialic acid axis promotes antitumor immune responses in preclinical models of glioblastoma. Sci. Transl. Med. 15, eadf5302 (2023).

50. G. Cyster, D. M. Shotton, A. F. Williams, The dimensions of the T lymphocyte glycoprotein leukosialin and identification of linear protein epitopes that can be modified by glycosylation. EMBO J. 10, 893–902 (1991).

51. A. Baudino, Y. Shinohara, F. Nimmerjahn, J.-I. Furukawa, M. Nakata, E. Martínez-Soria, F. Petry, J. V. Ravetch, S.-I. Nishimura, S. Izui, Crucial role of aspartic acid at position 265 in the CH2 domain for murine IgG2a and IgG2b Fc-associated effector functions. J. Immunol. 181, 6664–6669 (2008).

52. R. A. Franklin, W. Liao, A. Sarkar, M. V. Kim, M. R. Bivona, K. Liu, E. G. Pamer, M. O. Li, The cellular and molecular origin of tumor-associated macrophages. Science 344, 921–925 (2014).

53. A. Robinson, C. Z. Han, C. K. Glass, J. W. Pollard, Monocyte Regulation in Homeostasis and Malignancy. Trends Immunol. 42, 104–119 (2021).

54. S. Cheng, Z. Li, R. Gao, B. Xing, Y. Gao, Y. Yang, S. Qin, L. Zhang, H. Ouyang, P. Du, L. Jiang, B. Zhang, Y. Yang, X. Wang, X. Ren, J.-X. Bei, X. Hu, Z. Bu, J. Ji, Z. Zhang, A pan-cancer single-cell transcriptional atlas of tumor infiltrating myeloid cells. Cell 184, 792–809.e23 (2021).

55. A. Reiss, M. G. Angelos, E. C. Dees, Y. Yuan, N. T. Ueno, P. R. Pohlmann, M. L. Johnson, J. Chao, O. Shestova, J. S. Serody, M. Schmierer, M. Kremp, M. Ball, R. Qureshi, B. H. Schott, P. Sonawane, S. C. DeLong, M. Christiano, R. F. Swaby, S. Abramson, K. Locke, D. Barton, E. Kennedy, S. Gill, D. Cushing, M. Klichinsky, T. Condamine, Y. Abdou, CAR-macrophage therapy for HER2-overexpressing advanced solid tumors: a phase 1 trial. Nat. Med., doi: 10.1038/s41591-025-03495-z (2025).

56. T. Yamauchi, K. Takenaka, S. Urata, T. Shima, Y. Kikushige, T. Tokuyama, C. Iwamoto, M. Nishihara, H. Iwasaki, T. Miyamoto, N. Honma, M. Nakao, T. Matozaki, K. Akashi, Polymorphic Sirpa is the genetic determinant for NOD-based mouse lines to achieve efficient human cell engraftment. Blood 121, 1316–1325 (2013).

57. A. Wang, Y. Hou, J. Zak, Q. Zheng, K. A. McCord, M. Wu, D. Zhang, S. Chung, Y. Shi, J. Ye, Y. Zhao, S. Hajjar, I. A. Wilson, J. C. Paulson, J. R. Teijaro, X. Zhou, K. B. Sharpless, M. S. Macauley, P. Wu, Reshaping the tumor microenvironment by degrading glycoimmune checkpoints Siglec-7 and −9, bioRxivorg (2024). 10.1101/2024.10.11.617879.

58. D. L. Savage, S. L. Kimzey, S. K. Bromley, K. G. Johnson, M. L. Dustin, J. M. Green, Polar redistribution of the sialoglycoprotein CD43: implications for T cell function. J. Immunol. 168, 3740– 3746 (2002).

59. A. Wu, X. Wang, Y. Huang, Y. Zhang, S. Su, H. Shou, H. Wang, J. Zhang, B. Wang, Targeted glycan degradation potentiates cellular immunotherapy for solid tumors. Proc. Natl. Acad. Sci. U. S. A. 120, e2300366120 (2023).

60. A. Stanczak, N. Rodrigues Mantuano, N. Kirchhammer, D. E. Sanin, F. Jacob, R. Coelho, A. V. Everest-Dass, J. Wang, M. P. Trefny, G. Monaco, A. Bärenwaldt, M. A. Gray, A. Petrone, A. S. Kashyap, K. Glatz, B. Kasenda, K. Normington, J. Broderick, L. Peng, O. M. T. Pearce, E. L. Pearce, C. R. Bertozzi, A. Zippelius, H. Läubli, Targeting cancer glycosylation repolarizes tumor-associated macrophages allowing effective immune checkpoint blockade. Sci. Transl. Med. 14, eabj1270 (2022).

61. M. A. Gray, M. A. Stanczak, N. R. Mantuano, H. Xiao, J. F. A. Pijnenborg, S. A. Malaker, C. L. Miller, P. A. Weidenbacher, J. T. Tanzo, G. Ahn, E. C. Woods, H. Läubli, C. R. Bertozzi, Targeted glycan degradation potentiates the anticancer immune response in vivo. Nat. Chem. Biol. 16, 1376–1384 (2020).

62. A. Pinello, M. C. Canver, M. D. Hoban, S. H. Orkin, D. B. Kohn, D. E. Bauer, G.-C. Yuan, Analyzing CRISPR genome-editing experiments with CRISPResso. Nat. Biotechnol. 34, 695–697 (2016).

63. A. Clement, H. Rees, M. C. Canver, J. M. Gehrke, R. Farouni, J. Y. Hsu, M. A. Cole, D. R. Liu, J. K. Joung, D. E. Bauer, L. Pinello, CRISPResso2 provides accurate and rapid genome editing sequence analysis. Nat. Biotechnol. 37, 224–226 (2019).

64. J. G. Doench, N. Fusi, M. Sullender, M. Hegde, E. W. Vaimberg, K. F. Donovan, I. Smith, Z. Tothova, C. Wilen, R. Orchard, H. W. Virgin, J. Listgarten, D. E. Root, Optimized sgRNA design to maximize activity and minimize off-target effects of CRISPR-Cas9. Nat. Biotechnol. 34, 184–191 (2016).

65. R. A. Flynn, K. Pedram, S. A. Malaker, P. J. Batista, B. A. H. Smith, A. G. Johnson, B. M. George, K. Majzoub, P. W. Villalta, J. E. Carette, C. R. Bertozzi, Small RNAs are modified with N-glycans and displayed on the surface of living cells. Cell 184, 3109–3124.e22 (2021).

66. F. A. Wolf, P. Angerer, F. J. Theis, SCANPY: large-scale single-cell gene expression data analysis. Genome Biol. 19, 15 (2018).

67. S. L. Wolock, R. Lopez, A. M. Klein, Scrublet: Computational identification of cell Doublets in Single-cell transcriptomic data. Cell Syst. 8, 281–291.e9 (2019).

68. R. A. Amezquita, A. T. L. Lun, E. Becht, V. J. Carey, L. N. Carpp, L. Geistlinger, F. Marini, K. Rue-Albrecht, D. Risso, C. Soneson, L. Waldron, H. Pagès, M. L. Smith, W. Huber, M. Morgan, R. Gottardo, S. C. Hicks, Orchestrating single-cell analysis with Bioconductor. Nat. Methods 17, 137–145 (2020).

69. A. L. Luke Zappia, Zellkonverter (Bioconductor, 2020; 10.18129/B9.BIOC.ZELLKONVERTER).

