## Supplemental Figures for "Sialylated CD43 is a glyco-immune checkpoint for macrophage phagocytosis"

**The PDF file includes:**

Materials and Methods
Figs. S1 to S11
Tables S1 to S11

#### Materials and Methods

##### Isolation of human monocytes and differentiation into human macrophages

Peripheral blood mononuclear cells (PBMCs) were isolated using Ficoll gradient separation of whole blood from healthy donors. Monocytes were isolated from PBMCs using negative selection using the EasySep Human Monocyte Enrichment Kit (Stemcell Technologies). Following isolation, monocytes were resuspended in RPMI 1640 medium with 10% fetal bovine serums, penicillin, streptomycin, glutamine and 50 ng/mL of M-CSF (Peprotech). Monocytes were allowed to adhere to the tissue culture plate and media was exchanged every 2-3 days. Following differentiation, the macrophages were either stimulated with interferon-γ (Peprotech) or used directly for co-culture assays.

##### In vitro macrophage co-culture assays

Macrophages were plated in 24-well plates with 200,000 macrophages per well and co-cultured with leukemia cell lines at an effector to target ratio of 2:1. Control or knockout leukemia cell lines were engineered to express either BFP or RFP657 and 1:1 mixes of control and knockout lines with different fluorophores were added to macrophages. Following 18 hours of co-culture with or without macrophages, leukemia cells were harvested, washed and stained for CD11b-FITC as a macrophage marker prior to performing flow cytometry to distinguish the relative ratios of BFP or RFP expressing cells remaining in the culture. After staining with antibodies targeting CD11b, cells were resuspended in PBS with 2% FBS and 5 mM EDTA along with a fixed proportion of counting beads (Biolegend 424902) to normalize event acquisition across conditions. Samples were acquired on the Beckman Coulter Cytoflex instrument and analyzed using FlowJo software.

##### Mouse bone marrow-derived macrophage differentiation

Total bone marrow cells were flushed from the femurs and tibias of C57BL/6 mice with ice cold DMEM complete medium (DMEM + 10% FBS + 0.5% P/S). Cells were strained through a nylon mesh filter before red blood cell lysis with ACK buffer. Mouse bone marrow cells were then counted, resuspended in complete DMEM + 25 ng/mL mouse M-CSF, and plated in 15 cm dishes. Media was replenished on day 3 before adherent bone marrow-derived macrophages were lifted with 2.5 mM EDTA in PBS and plated for downstream phagocytosis experiments.

##### Cell culture

MV411, MOLM13, and HEL cell lines were cultured in RPMI 1640 (Sigma) supplemented with 10% fetal bovine serum and antibiotics. The identity of each cell line was confirmed by STR genotyping. All cell lines were negative for *Mycoplasma*.

##### Generation of CRISPR edited lines

For CRISPR knockout generation, cell lines stably expressed *S. pyogenes* Cas9 or *enCas12a* with a blasticidin selection cassette. Cell lines were infected with lentivirus containing sgRNAs targeting the gene of interest and a puromycin resistance cassette. Cell lines were selected with the 2-4 ug/mL of puromycin for 48 hours and cultured for approximately one week before characterization of the knockout phenotype using western blotting, flow cytometry, or sequencing of the sgRNA targeting DNA locus. CRISPRESSO2 (*62*, *63*) was used to quantify editing at the DNA locus targeted by the sgRNA of interest.

##### Cas9/RNP Editing of Monocytes

sgRNAs targeting genes of interest or a non-targeting control were synthesized as modified RNAs from Synthego. sgRNAs were precomplexed in a 1:1 ratio with Alt-R-V3 Cas9 from IDT at 37C for 15 minutes. Cas9/RNP complexes were electroporated into healthy donor derived monocytes immediately after isolation using the Lonza Nucleofector 4D and a P3 Primary Cell Nucleofector Kit (V4XP-3024). Following electroporation, monocytes were cultured in RPMI 1640 media with 10% FBS and 1% penicillin/streptomycin/glutamine supplemented with 50 ng/mL of M-CSF to induce macrophage differentiation. After 5-7 days, macrophages were harvested and flow cytometry was performed to validate knockout of the genes of interest.

**Genome-wide CRISPR co-culture screening**

For the genome-scale in vitro CRISPR screens, we used a MV411 cell line stably expressing enCas12a or the MOLM13 cell line stably expressing *S. pyogenes* Cas9. Cas9 or enCas12a activity was verified to be greater than 70%. MV411 expressing enCas12a was infected with set C and set D of the enCas12a Humagne library and the MOLM13 Cas9 expressing cell line was infected with the Brunello library. All pooled infections were performed to achieve infection rates of 15-30% to ensure single copy sgRNA infection at sufficient cell numbers to yield >1000x coverage per sgRNA. Transduced cells were selected with 4 ug/mL of puromycin for 48 hours and aliquots were frozen following selection for an early time point control of library representation. Transduced and selected cells were expanded for 5-7 days prior to performing co-culture with macrophages for 18 hours. Cells were maintained in culture to achieve >2000x representation and co-cultures were performed to maintain 500x-1000x representation. Following co-culture, remaining leukemia cells were separated from the macrophages and harvested. Macrophages which are tightly adherent to the tissue culture dish were harvested using TrypLE and physical agitation. DNA was isolated using the Qiagen Blood Maxi Kit and sgRNAs were PCR amplified using Argon and Kermit P5 and P7 illumina primers, prior to sequencing on an Illumina HiSeq.

**Co-culture CRISPR screen analysis**

Guide sequences were demultiplexed and quantified using PoolQ 3.6.1. Read counts were library normalized per 1 million reads and log2 transformed with a pseudocount of one. Gene-targeting guides were z normalized by the control sgRNA distribution. Guide fold changes for the n top performing guides (4 for MOLM13 Cas9 Brunello library; 2 for MV411 enCas12Humagne library) were calculated as residuals fit to a natural cubic spline with 4 degrees of freedom. As previously described, significant depleted or enriched sgRNAs were identified using the STARS algorithm (*64*) considering the n top performing mapped guides per gene. Average p-values for genes in the library were calculated with the hypergeometric distribution; a control distribution for this calculation was created by grouping together n random control guides (n = 4 for Brunello; n = 2 for Humagne) into pseudogenes.

**Flow cytometry**

For flow cytometry of cell lines, cells were isolated and washed with PBS with 2% FBS and 5 mM EDTA prior to staining with the indicated fluorescently labeled antibodies at concentrations recommended by the manufacturer. Assays were performed with $10^5$-$10^6$ cells and analyses were acquired using a Beckman Coulter Cytoflex instrument and analyzed using FlowJo software.

**Lectin staining**

Flow cytometry-based lectin staining (Vicia Villosa Lectin, Anti-Peanut Agglutinin, Sambucus Nigra Lectin, Maackia Amurensis Lectin II) was performed on human leukemia cell lines and primary AML blasts. Approximately $10^5$-$10^6$ cells were isolated per each condition and washed with PBS with 0.5% BSA prior to staining at 4°C with VVA-FITC (1:1000, Vector Laboratories, catalog FL-1231-2), VVA-Biotin (1:500, Vector Laboratories, catalog B-1235-2), PNA-Biotin (1:500, Vector Laboratories, catalog BA-0074-.5), SNA-Biotin (1:400, Vector Laboratories, catalog B-1305-2), or MAL II-Biotin (1:200, catalog B-1265-1). Samples that were stained with biotinylated lectins were washed twice and stained with streptavidin-FITC (1:200) or streptavidin-BV421 (1:200) at 4°C.

**SIGLEC Fc staining**

Flow cytometry-based staining of human leukemia cell lines and primary AML blasts was adapted from previous literature (*46, 65*). Briefly, approximately $10^5$-$10^6$ cells cells were washed with 0.5% BSA, blocked with human Fc blocking solution (TruX, Biolegend catalog 422302) before incubation at 4°C with pre-complexes consisting of 1 ug/mL SIGLEC 7-Fc (R&D, catalog 1138-SL) or 1 ug/mL SIGLEC 9-Fc (R&D, catalog 1139-SL) and goat anti-human IgG AlexaFluor488 (Jackson Immunoresearch, 109-545-003). Pre-complexes were formed for 1 hr at 4°C.

**Fluorescent live cell imaging**

M-CSF differentiated human macrophages were plated in imaging-compatible 24 well flat-bottom plates (Ibidi) and either unstimulated or stimulated with 20 ng/mL IFN**γ** approximately 24 hours prior to co-culture with leukemia cells. $GFP^{+}$ MV411 leukemia cells were labeled with pHrodo Red AM (Invitrogen) per the manufacturer's instructions, opsonized with isotype (mouse IgG1), anti-CD43 (MEM59), or anti-CD47 (MIAP410), and added to unstimulated or IFN**γ**-stimulated human macrophages. Live cell imaging was performed directly in the 24 well flat-bottom plates on the Mica microscope (Leica Microsystems). Images were processed in Fiji.

**Bulk RNA-sequencing of human macrophages**

Bulk RNA-sequencing analysis was performed as described in (citation). Briefly, Illumina adapter sequences were trimmed using Trimmomatic (v0.36) and pre-/post-trimming quality control was done with FastQC (v0.11.7). Reads were quantified by pseudoalignment to hg38 using Kallisto (v0.46.0), and RNA-seq gene counts were quantified using the tximport package (v1.24.0) in R. From raw counts, a pseudocount of one was added to every data point to prevent zero values in a later log2 transformation step. The data was TPM normalized to account for differences in transcript length and sequencing depth. Afterwards, the data was log2 transformed to ensure normality. The three biological replicates for each condition were then mean collapsed resulting in a gene expression value. To generate heatmaps of gene expression data, the gene expression values for the SIGLEC genes were plotted across the three interferon conditions using the R package ComplexHeatmap (v2.18.0). Hierarchical clustering was determined using the package's default complete linkage method with the euclidean distance.

**Single cell RNA-sequencing analysis**

The van Galen et. al., 2019 scRNA-Seq dataset was extracted from the Curated Cancer Cell Atlas, including cell type annotations, with the data converted to h5ad format via Scanpy(*66*). The data was then filtered to remove cells composed of > 10% mitochondrial genes and doublets with Scrublet(*67*). The data was normalized via the shifted logarithm approach, implemented by the usage of the Scanpy preprocessing 'normalize_total' and 'log1p' functions. The data was then loaded into a SingleCellExperiment(*68*) object using zellkonverter(*69*). Cells were then Z-scored for their normalized expression of genes of interest and grouped by their cell type and cell subtype, with cells originating from post-treatment patient samples being removed. The mean Z-score and the percentage of cells expressing each gene of interest for each cell type/subtype group was then calculated and plotted.

***In vivo* models of leukemia**

MV411 leukemias (sgCtrl, sgC1GALT1, sgCD43) were infected with mouse CD8**α** lentivirus (CD8a-pLX313) and confirmed to express high levels of mouse CD8**α** by flow cytometry prior to use in experiments. 0.5x106 MV411 leukemias were injected intravenously into sublethally irradiated (225 Gy) NOD $Prkdc^{scid}$ $Il2rg^{null}$ (NSG) mice. Beginning on day 5 after injection, mice were injected with anti-mouse CD8 (clone 53-6.7) intraperitoneally until mice reached their endpoint.

**Glycopeptide Ranking**

To identify and prioritize genes within the O-linked glycosylation pathway, we first calculated a "Macrophage Enrichment Score" by summing the absolute max-normalized gene log-fold changes from the macrophage arms of the "IFN-high vs Isotype" comparisons in the non-ADCP IFNγ-stimulated

MV411 enCas12 Humagne and MOLM13 Cas9 Brunello screens. Cell line RNA-sequencing for MOLM13 (ACH-000362) and MV411 (ACH-000045) was collected from the Cancer Cell Line Encyclopedia (CCLE). Data was subsetted to genes coding for proteins classified as "membrane" or "membrane and secreted isoforms" (Human Protein Atlas) and as O-linked glycosylated peptides (*35, 36*). Numbers of glycosylation sites per protein were derived from GlyCosmos (*37*); only peptides with a recorded number of sites were maintained in the analysis.

**Culturing of AML patient samples**

Peripheral blood mononuclear cells were isolated from peripheral blood from patients with acute myeloid leukemia and peripheral blood blast count >20% who provided informed consent prior to inclusion in the study. Isolated cells were cultured in short-term expansion media consisting of StemSpan II (Stemcell Technologies Cat No. 9655), CC100 (Stemcell technologies Cat No. 2690), 100 ng/mL human TPO (Peprotech Cat No. 10773-602), L-glutamine (Life Technologies Cat No. 25030081), and Penicillin-Streptomycin (Life Technologies Cat No. 15140163). Isolated primary AML cells were cultured either in the presence or absence of macrophages.

**Cell binding avidity assays**

$10^7$ IFN**γ**-stimulated human macrophages were attached to poly-L-lysine coated chips to form a monolayer >80% confluency. Control or CD43-deficient MV411 leukemia cells were labeled with CellTrace Far Red dye (Thermo Fisher, catalog C34564) and 200 uL of cells were added to the human macrophage monolayers at a concentration of $10^6$/mL. After a 5 minute incubation period, increasing amounts of force was applied with the z-Movi Cell Avidity Analyzer (Lumicks) and cell detachment was quantified. Analysis was performed using Ocean software.

**Western blotting**

Cell pellets were lysed in RIPA buffer supplemented with 5 mM EDTA and protease inhibitor cocktail (Thermo Scientific). Cell pellets were incubated and rotated on a rocker at 4C for 30 minutes. Total protein amounts were quantified by Bicinchoninic Acid (BCA) assay. Samples were denatured in sample buffer containing DTT and boiled at 95C for 5-10 minutes. Approximately 30-40 ug of protein was loaded into each well of 4-12% gradient Bis-Tris gels and resolved by SDS-PAGE. Gels were transferred onto PVDF membranes via wet transfer for 2 hours at 4C. Membranes were blotted overnight with primary antibodies directed against sialylated CD43, total CD43, and/or GAPDH. Membranes were washed, stained with fluorescently tagged secondary antibodies (Licor), and visualized on a Licor machine.

**Reagent table**

CRISPR sgRNA sequences

| **Gene** | **Cas9 sgRNA sequences** |
|---|---|
| SPN | GATCCCACACCGTGACAGG |
| C1GALT1 | TCATACTAGACAATTTGAGG |
| C1GALT1C1 | CAATGATAGCAAACGTAGTG |
| SLC35A2 | TAGAGATGGCAACATACTGG |
| SLC39A9 | TATGTTGGTGGGATGTTACG |
| TAP1 | CCCAGAGTACAGAAGGCTGT |

| | |
|---|---|
| NLRC5 | GTGCTCTAGGATGTTGGTCA |
| TAPBP | GCCGCTGGCCCATTTCGCAG |
| CMAS | AGAACATTAAGCACCTGGCG |
| SLC35A1 | TTCTGTGATACACACGGCTG |
| LILRB1 | ATTCCCTCCTGAGTTCACCA |
| LILRB2 | CCGTCACCCTCAGTTGTCAG |
| SIGLEC7_9 | TCTGACCTGCTCTGTGCCCT |
| CD47 | ACTCTTATCCATCTTCAAAG |
| MGAT1 | AAAGTACTCGAAGAAGTCCG |
| MGAT1 | CATCGCCTCCTACGGCAGCG |

| **Construct** | **enCas12a sgRNA sequence** |
|---|---|
| CD43_Ctrl | GACTTCCATTGGTGCCAGCACTGTAATTTCTACTGTCGTAGATACGCTAGAGACCTGGGGTCCACTTAATTTCTACTATCGTAGATTGAAGTTCAATACATCGCGTCTTAAATTTCTACTCTAGTAGATACACACTAGAACTTCGCCATGAG |
| CD43_C1GALT1 | GACTTCCATTGGTGCCAGCACTGTAATTTCTACTGTCGTAGATACGCTAGAGACCTGGGGTCCACTTAATTTCTACTATCGTAGATTCCCAACAAAATACTAAATAGCTAAATTTCTACTCTAGTAGATCATAAGGCTTAAATCTTCTCCCA |
| CD43_C1GALT1C1 | GACTTCCATTGGTGCCAGCACTGTAATTTCTACTGTCGTAGATACGCTAGAGACCTGGGGTCCACTTAATTTCTACTATCGTAGATTATCTAGGCCACACTATAAAATCAAATTTCTACTCTAGTAGATGTCCAAGTCTCCTTTACTGCAGC |
| C1GALT1_Ctrl | TCCCAACAAAATACTAAATAGCTTAATTTCTACTGTCGTAGATCATAAGGCTTAAATCTTCTCCCATAATTTCTACTATCGTAGATTGAAGTTCAATACATCGCGTCTTAAATTTCTACTCTAGTAGATACACACTAGAACTTCGCCATGAG |
| C1GALT1C1_Ctrl | TATCTAGGCCACACTATAAAATCTAATTTCTACTGTCGTAGATGTCCAAGTCTCCTTTACTGCAGCTAATTTCTACTATCGTAGATTGAAGTTCAATACATCGCGTCTTAAATTTCTACTCTAGTAGATACACACTAGAACTTCGCCATGAG |
| Ctrl_Ctrl | ACACCATACCCGGGACCACTGCTTAATTTCTACTGTCGTAGATCCATTCAGGTCGATCGGATTGGCTAATTTCTACTATCGTAGATGCTACCAAGGGTGTTTGTCCCTGAAATTTCTACTCTAGTAGATCTCCAGGCGAGCGTGTTCGCTCC |

**Antibodies**

The following antibodies were used for flow cytometry: anti-human CD11b (clone M1/70, Biolegend), anti-human CD43 (MEM59, Thermo Fisher), anti-human CD43 (10G7, Biolegend), anti-mouse CD8a (53-6.7, Biolegend), anti-human CD34 (clone 581, Biolegend). The following antibodies were used for western blot analysis: anti-human MGAT1 (clone EPR1247, Abcam), anti-human sialylated CD43 (clone MEM59, Thermo Fisher), anti-human CD43 (clone PA5-80067, Thermo Fisher), anti-PTPN6 (clone E1U6R, Cell Signaling).

**Fig. S1.**

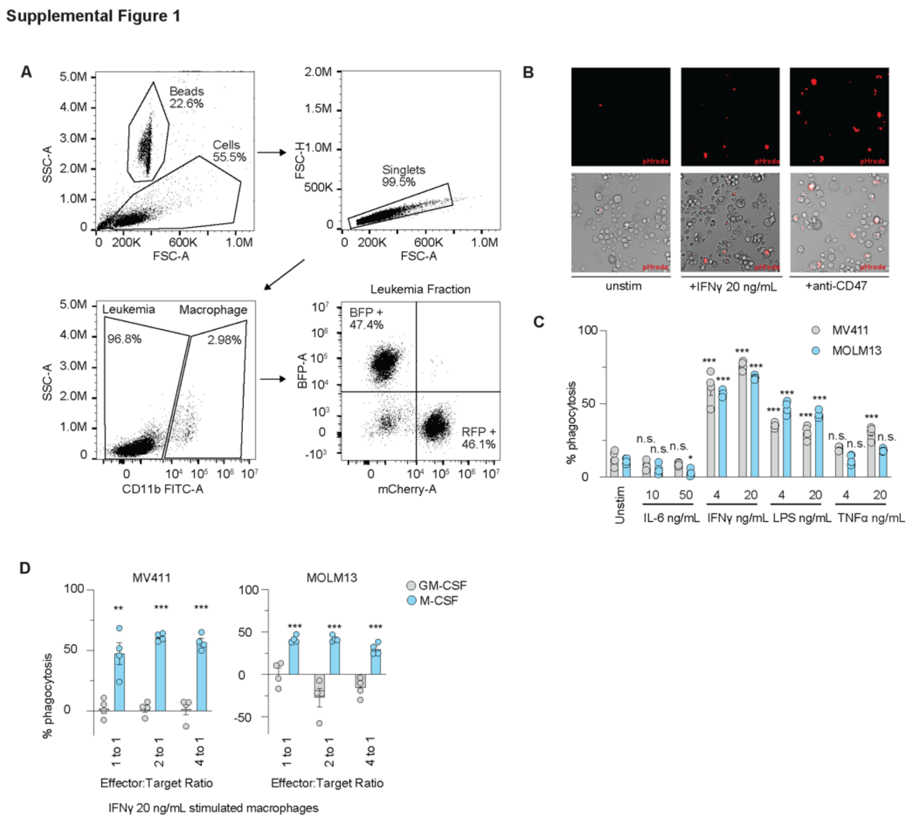


**Fig. S1:Establishment of a human macrophage in vitro co-culture assay a,** Flow cytometry gating strategy to quantify and assess phagocytosis. Macrophages are CD11b-FITC positive and leukemia cells express RFP or BFP. Gating strategy is shown. **b,** Live cell microscopy of pHrodo stained leukemia cells co-cultured with macrophages that are unstimulated, stimulated with IFN**γ**, or following leukemia treatment with Anti-CD47. **c**, Quantification of phagocytosis with MCSF differentiated macrophages following treatment with various cytokines. **d,** Assessment of percent of phagocytosis seen with M-CSF or GM-CSF differentiated macrophages stimulated with IFN**γ** at a range of effector to target ratios. Data in **c** were analyzed by two-way ANOVA, and data in **d** were analyzed with unpaired, two-sided Student's *t*-test, * $p<0.05$, ** $p<0.01$, *** $p<0.001$.

#### Fig. S2.

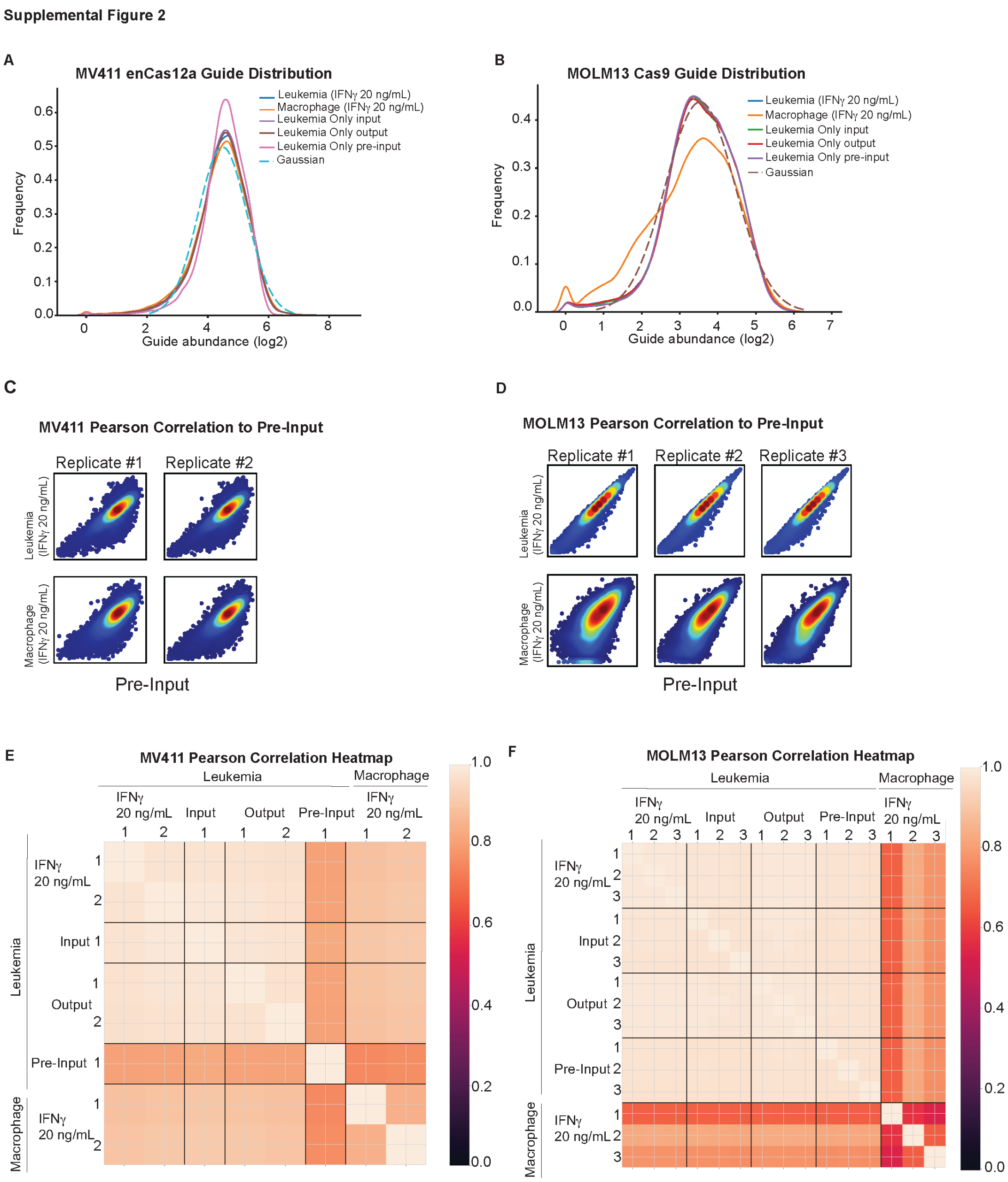


**Fig. S2: Genetic screen quality metrics for antibody independent cellular phagocytosis screens. a** and **b,** Plot of the abundance of sgRNAs in leukemic cells in the indicated conditions compared to a gaussian distribution of sgRNAs in MV411 and MOLM13. **b,** and **c,** Net replicate Pearson correlation of each individual replicate versus pre-input representing the initial distribution of the sgRNA library in MV411 and MOLM13. **d,** and **e,** Pearson correlation heatmap of replicates in the leukemia fraction and macrophage fraction versus one another in the MV411 and MOLM13 antibody independent cellular phagocytosis screens, respectively.

**Fig.S3.**

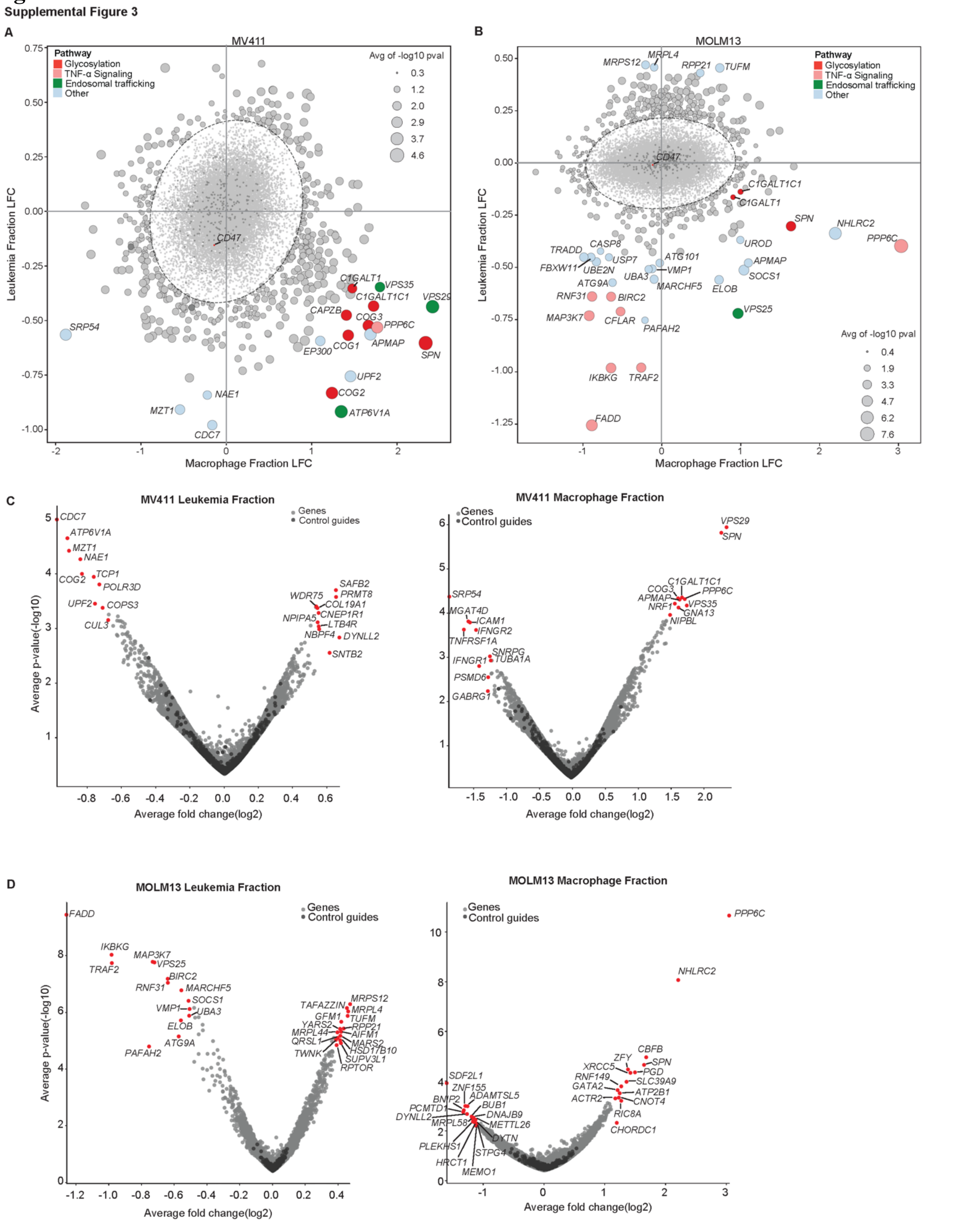


**Fig. S3: Top hits that modulate antibody independent cellular phagocytosis a,** Scatter plot of log fold change (LFC) of enrichment or depletion of sgRNAs in the leukemia fraction versus leukemia only (*y*-

axis) and macrophage fraction versus leukemia only (*x*-axis) for MV411. **b,** Scatter plot of LFC values of enrichment or depletion of sgRNAs in the leukemia fraction versus leukemia only (*y*-axis) versus macrophage fraction versus leukemia only (*x*-axis). Circle sizes for points in **a,** and **b,** are scaled by average log10 p-value. **c,** Volcano plot of LFC of MV411 leukemia fraction or MV411 macrophage fraction (*x*-axis) versus the average -log10(p-value) (*y*-axis). **d,** Volcano plot of LFC of MV411 leukemia fraction or macrophage fraction (*x*-axis) versus the average -log10(p-value) (*y*-axis).

**Fig. S4.**

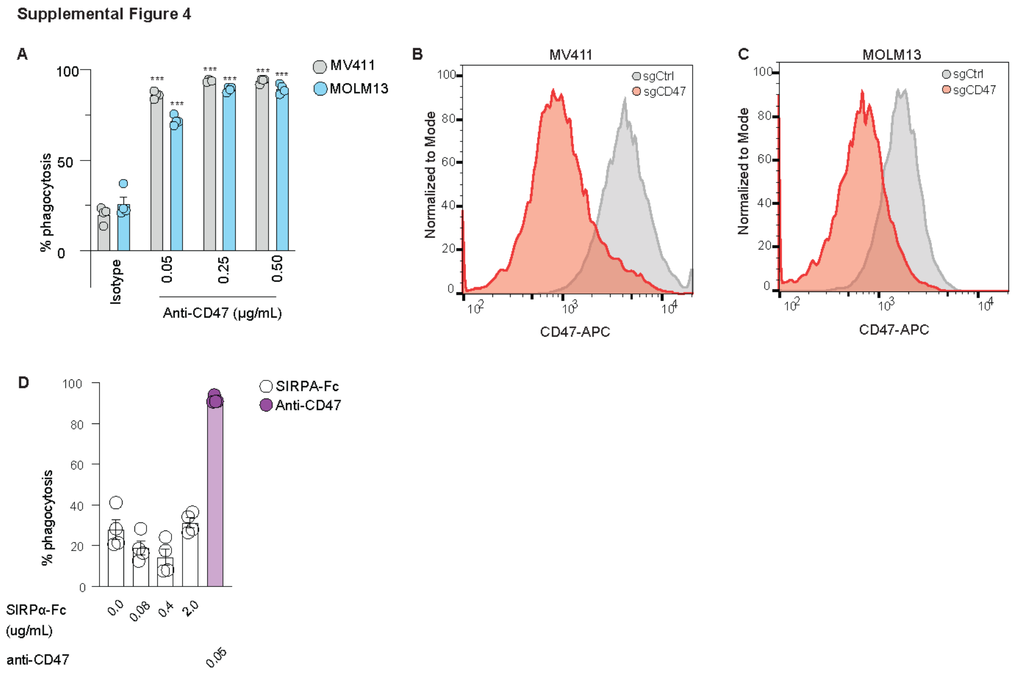


**Fig. S4: Impact of targeting CD47 with antibodies or genetic deletion on macrophage phagocytosis.** **a,** Percent of leukemia cells phagocytosed following treatment with isotype control or varying doses of anti-CD47 (MIAP410) antibody. Data show the mean ± s.e.m. of four technical replicates and are representative of two different experiments. **b,** Flow cytometry of CD47 expression in MV411 Cas9 cells transduced with sgRNAs targeting CD47 or a control locus. **c**, Flow cytometry of CD47 expression in MV411 Cas9 cells transduced with sgRNAs targeting CD47 or a control locus. **d,** Percent of leukemia cells phagocytosed following treatment with SIRPα-Fc or Anti-CD47 (MIAP410) antibody. Data show the mean ± s.e.m. of four technical replicates and are representative of two different experiments. Data in **a** were analyzed by two-way ANOVA, *** $p<0.001$.

**Fig. S5.**

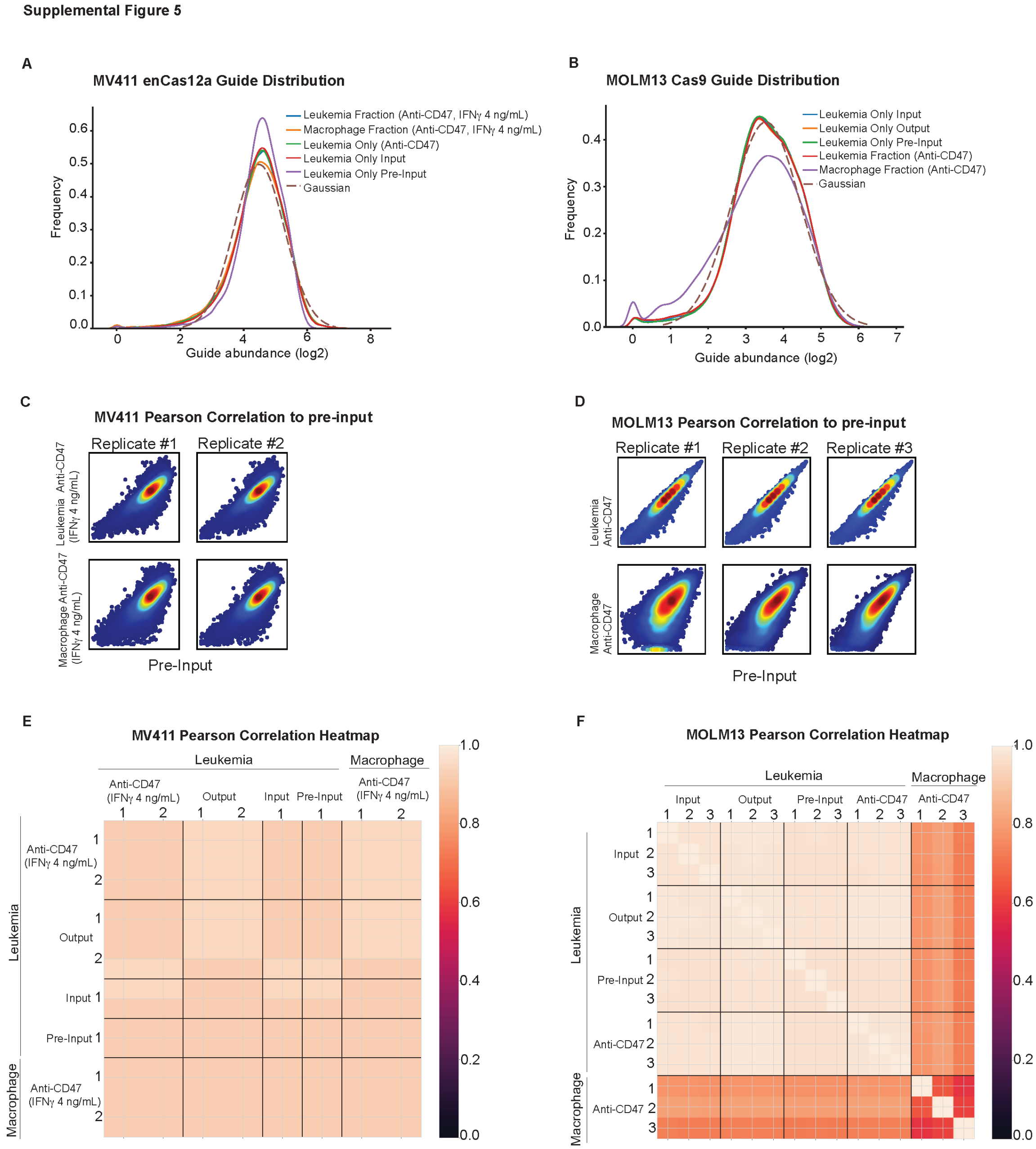


**Fig. S5: Genetic screen quality metrics for antibody dependent cellular phagocytosis screens. a** and **b,** Plot of the abundance of sgRNAs in leukemic cells in the indicated conditions compared to a gaussian distribution of sgRNAs in MV411 and MOLM13 antibody-dependent cellular phagocytosis screens. **b,** and **c,** Net replicate Pearson correlation of each individual replicate versus pre-input representing the initial distribution of the sgRNA library in MV411 and MOLM13. **d** and **e,** Pearson correlation heatmap of all replicates in the leukemia fraction and macrophage fraction versus one another in the MV411 and MOLM13 antibody independent cellular phagocytosis screens, respectively.

#### Fig. S6.

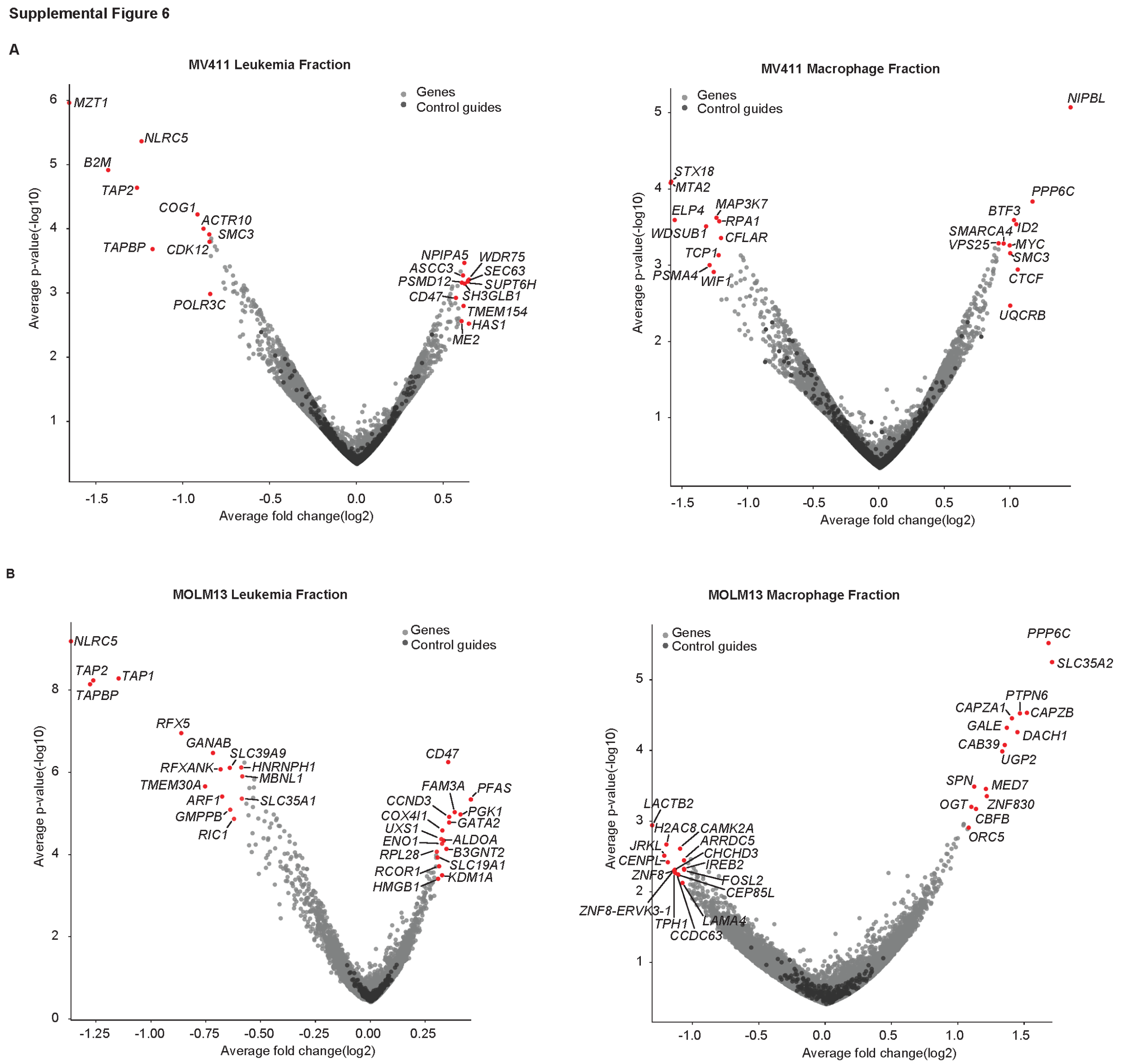


**Fig. S6:Top hits that modulate antibody dependent cellular phagocytosis. a,** Volcano plot of LFC of MV411 leukemia fraction or MV411 macrophage fraction (*x*-axis) versus the average -log10(p-value) (*y*-axis). **b,** Volcano plot of LFC of MV411 leukemia fraction or macrophage fraction (*x*-axis) versus the average -log10(p-value) (*y*-axis).

### Fig. S7.

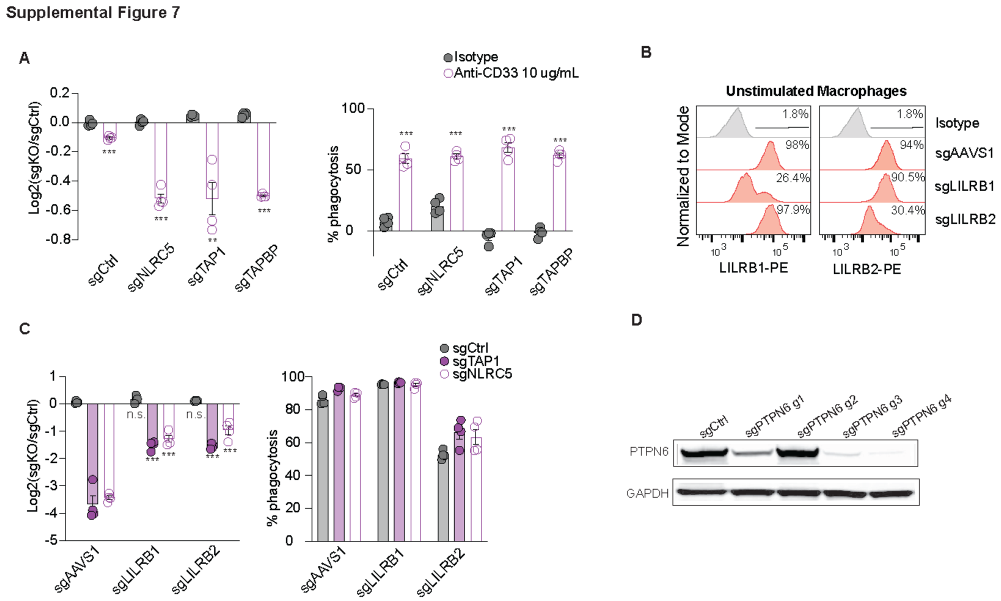


**Fig. S7: Validation of genetic factors that modulate antibody dependent cellular phagocytosis. a,** Ratio of phagocytosis of MV411 leukemia cells with MHC-I loss versus control leukemia in co-culture. Leukemia cells were treated with either anti-isotype or anti-CD33. Percent of leukemia cells is quantified in each condition. **b,** flow cytometry of macrophages following editing with Cas9-RNP with sgRNAs targeting AAVS1, LILRB1, or LILRB2. Expression of LILRB1 or LILRB2 is shown. Percent of macrophages positive for LILRB1 or LILRB2 expression is quantified. **c,** Quantification of the ratio of phagocytosis of MV411 leukemia with MHC-I loss versus control leukemia in co-culture with sgAAVS1, sgLILRB1, or sgLILRB2 edited macrophages. Percent of leukemia cells phagocytosed is quantified in each macrophage background. **d,** Western blotting for PTPN6 in control MV411 leukemia or in leukemic cells transduced with sgRNAs targeting PTPN6. Data in **a,c** were analyzed with unpaired, two-sided Student's *t*-test, * p<0.05, ** p<0.01, *** p<0.001.

#### Fig. S8.

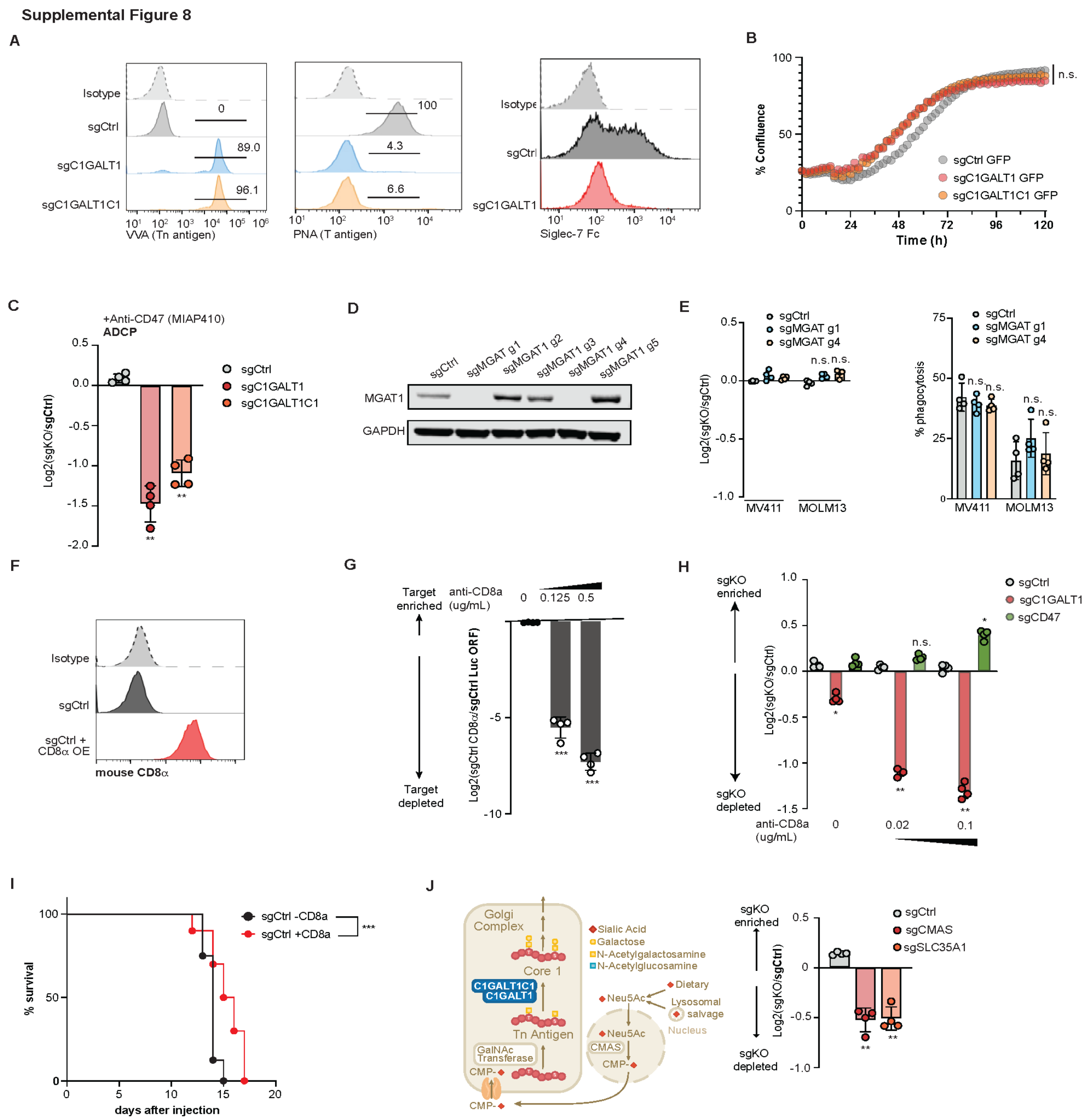


**Fig. S8: Loss of O-glycosylation and cell surface sialylation enhances phagocytosis *in vitro* and *in vivo*.** **a**, VVA, PNA, and SIGLEC 7-Fc staining of control, C1GALT1-deficient, or C1GALT1C1-deficient MV411 leukemias **b**, Incucyte-based growth curves for control, C1GALT1-deficient, or C1GALT1C1-deficient MV411 leukemias **c**, Bar graph of relative antibody-dependent cellular phagocytosis of control, C1GALT1-deficient, or C1GALT1C1-deficient MV411 leukemias by unstimulated macrophages after addition of anti-CD47 antibodies **d**, Representative western blots for MGAT1 expression after editing MV411 cells with various MGAT1 sgRNAs. **e**, Relative and absolute phagocytosis of control or MGAT1-deficient leukemias versus control leukemias after co-culture with IFN**γ**-stimulated macrophages **f**, Representative flow cytometry plots of mouse CD8**α** overexpression in MV411 leukemias **g**, Bar graphs of relative phagocytosis of control leukemias overexpressing mouse CD8**α** versus control leukemias overexpressing luciferase in the presence of various doses of anti-CD8**α** antibodies **h**, Bar graphs of relative phagocytosis of control, C1GALT1-deficient, or CD47-deficient leukemias cells overexpressing mouse CD8**α** relative to control leukemias overexpressing mouse CD8**α**

in the presence of varying doses of CD8**α** antibodies **i**, Survival of mice challenged with MV411 overexpressing mouse CD8**α** leukemias treated with isotype or anti-CD8**α** antibodies **j**, Schematic of sialic acid biosynthesis and relative phagocytosis of control, *CMAS*-deficient, or *SLC35A1*-deficient leukemias versus control MV411 leukemias after co-culture with IFN**γ**-stimulated macrophages. Data in **c**,**d**,**g**,**h**,**j** were analyzed with unpaired, two-sided Student's *t*-test, * $p<0.05$, ** $p<0.01$, *** $p<0.001$.

**Fig.S9.**

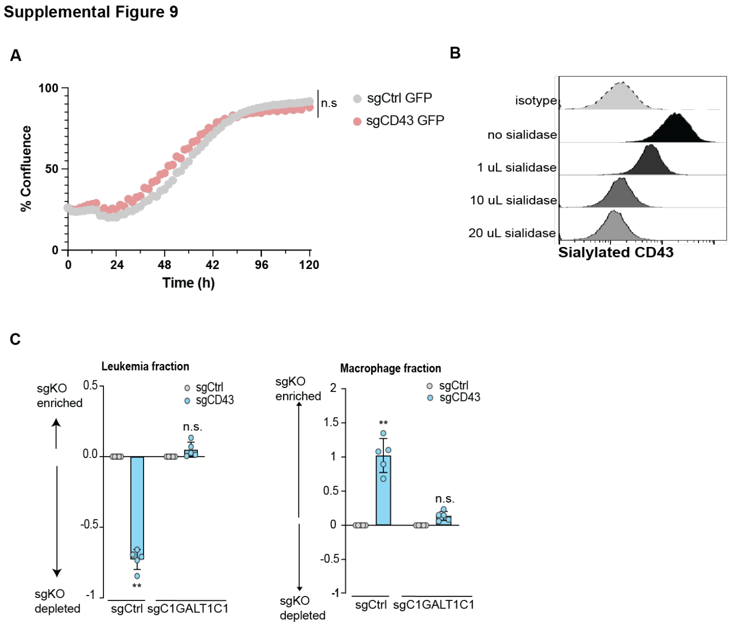


**Fig. S9: CD43 loss is epistatic with C1GALT1C1 loss. a,** Incucyte-based growth curves for control or CD43-deficient MV411 leukemias **b,** Representative flow cytometry plots of sialylated CD43 expression after treatment of leukemias cells with varying doses of *V. cholerae* sialidase **c,** Relative phagocytosis of CD43-deficient or dual C1GALT1C1- and CD43-deficient leukemias after co-culture with IFN**γ**-stimulated macrophages. Data in **c** were analyzed with unpaired, two-sided Student's *t*-test, ** p<0.01, n.s. not significant

**Fig. S10.**

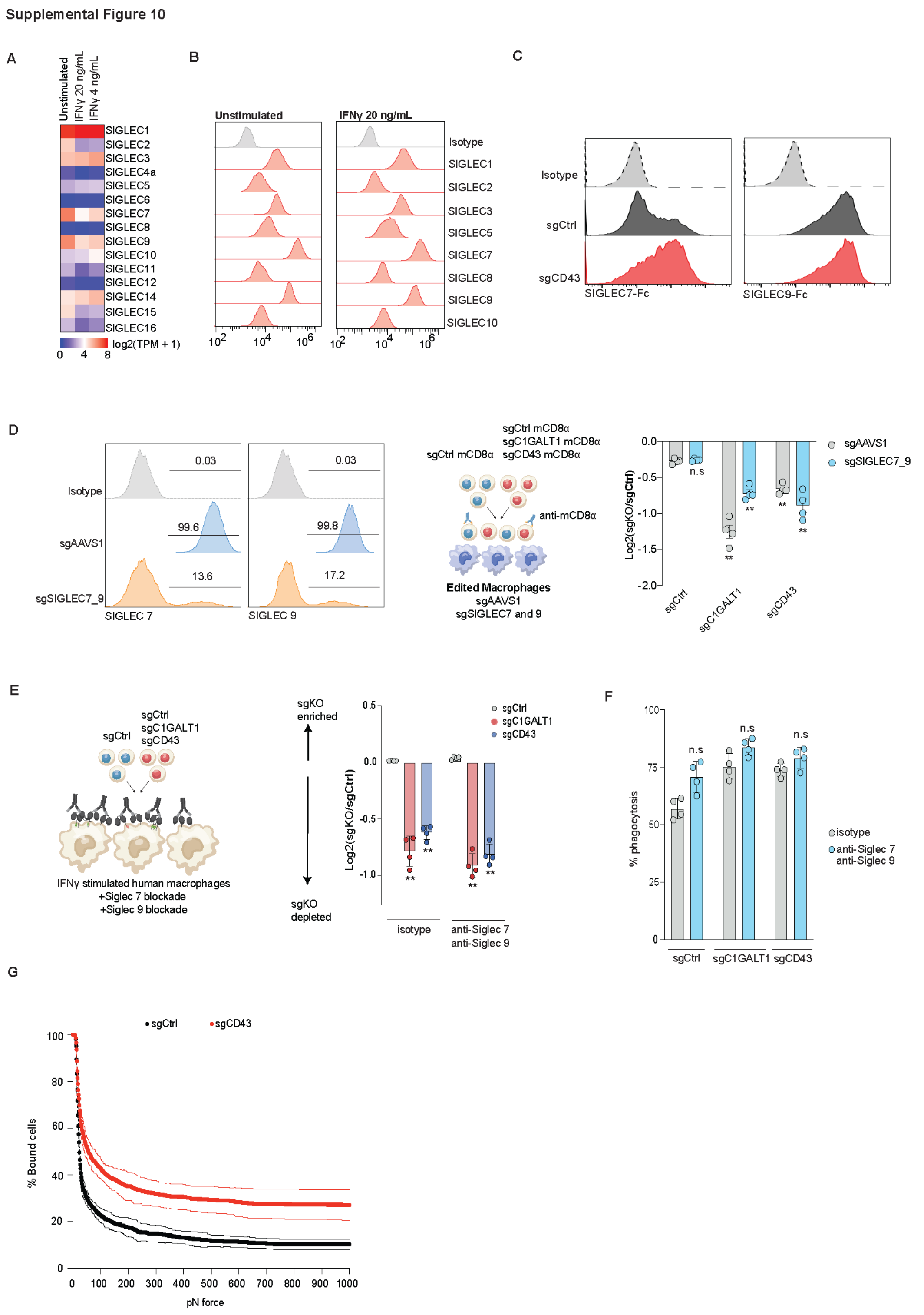


350 **Fig. S10: Loss or inhibition of SIGLEC7 and 9 do not modify phagocytosis of CD43 or C1GALT1C1 deficient leukemia cells. a,** RNA transcript levels of SIGLEC family members taken from bulk RNA-sequencing of unstimulated or IFN**γ**-stimulated human macrophages **b,** Representative flow

cytometry plots depicting SIGLEC family member expression in unstimulated or IFN**γ**-stimulated macrophages **c,** SIGLEC 7-Fc or SIGLEC 9-Fc staining on control or CD43-deficient MV411 leukemias **d,** Representative flow cytometry plots for SIGLEC 7 and SIGLEC 9 expression after genetic deletion of both SIGLEC 7 and SIGLEC 9 in human macrophages and relative phagocytosis of control, C1GALT1-deficient, or CD43-deficient leukemias by control or SIGLEC 7/9-deficient macrophages **e,** Relative phagocytosis of control, C1GALT1-deficient, or CD43-deficient leukemias by IFN**γ**-stimulated macrophages after addition of isotype or anti-SIGLEC 7/anti-SIGLEC 9 neutralizing antibodies **f,** absolute phagocytosis of control, C1GALT1-deficient, or CD43-deficient leukemias after addition of anti-SIGLEC 7/anti-SIGLEC 9 neutralizing antibodies **g,** Cell binding avidity measured by z-Movi of IFN**γ**-stimulated macrophages co-cultured with control or CD43-deficient MV411 leukemias (n=3). Data in **d,e** were analyzed by unpaired, two-sided Student's *t*-test, ** $p < 0.01$. Data in **g** are representative of three independent z-Movi chips.

**Fig. S11.**

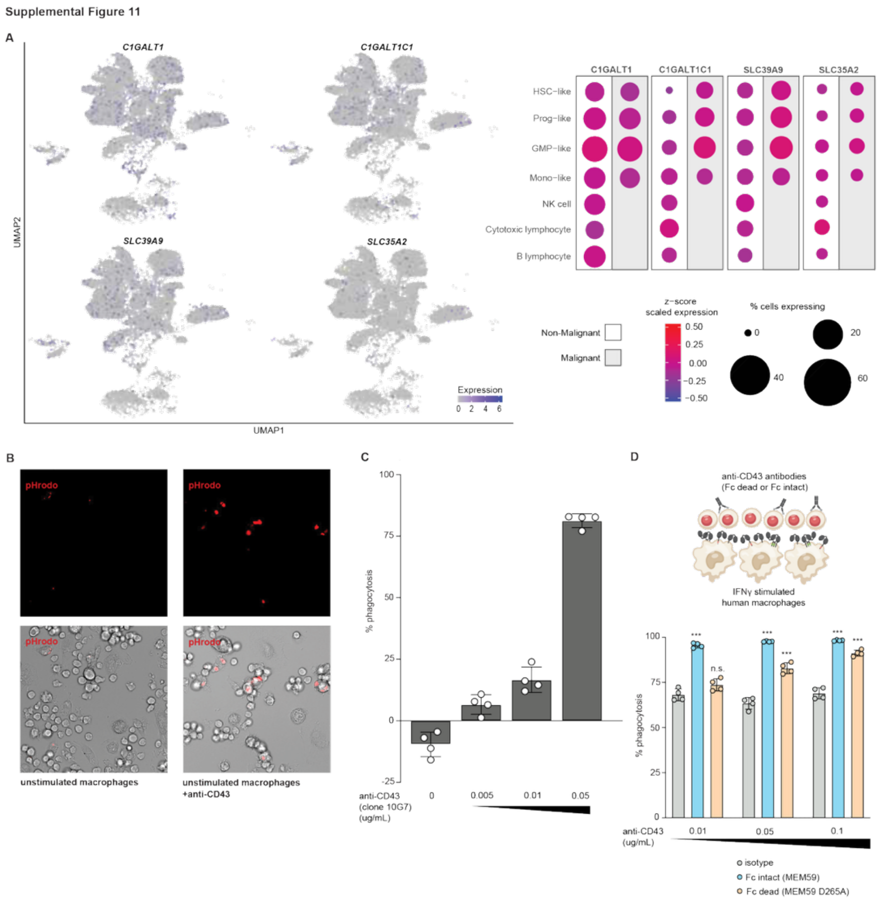


**Fig. S11: Expression O-glycan pathway genes and the impact of anti-CD43 targeting antibodies on phagocytosis of leukemia cells. a**, Expression of C1GALT1, C1GALT1C1, SLC39A9, and SLC35A2 across a UMAP projection of scRNA-seq profiles of bone marrow aspirate cells taken from healthy patients or AML patients. **b,** Representative fluorescence microscopy images of human macrophages co-cultured with pHrodo-labeled human leukemia cells in the presence or absence of anti-CD43 antibody (clone MEM59) **c,** Absolute phagocytosis of MV411 leukemia cells by unstimulated human macrophages after addition of various concentrations of anti-CD43 (clone 10G7) **d,** Absolute phagocytosis of MV411 leukemia cells by IFN**γ**-stimulated macrophages after addition of anti-CD43 antibodies in the presence or absence of Fc blockade. Data in **c,d** were analyzed by unpaired, two-sided Student's *t*-test, *p<0.05, ** p< 0.01.

**Table S1: CRISPR screen results for the leukemia fraction of antibody-independent cellular phagocytosis comparing IFNγ stimulated co-culture versus leukemia only in MV411**

**Table S2: CRISPR screen results for the leukemia fraction of antibody-independent cellular phagocytosis comparing IFNγ stimulated co-culture versus leukemia only in MOLM13**

**Table S3: CRISPR screen results for the macrophage fraction of antibody-independent cellular phagocytosis comparing IFNγ stimulated co-culture versus leukemia only in MV411**

**Table S4: CRISPR screen results for the macrophage fraction of antibody-independent cellular phagocytosis comparing IFNγ stimulated co-culture versus leukemia only in MOLM13**

**Table S5: CRISPR screen results for the leukemia fraction of antibody-dependent cellular phagocytosis comparing co-culture versus leukemia only in MV411**

**Table S6: CRISPR screen results for the leukemia fraction of antibody-dependent cellular phagocytosis comparing co-culture versus leukemia only in MOLM13**

**Table S7: CRISPR screen results for the macrophage fraction of antibody-dependent cellular phagocytosis comparing co-culture versus leukemia only in MV411**

**Table S8: CRISPR screen results for the macrophage fraction of antibody-dependent cellular phagocytosis comparing co-culture versus leukemia only in MOLM13**

**Table S9: Genome library sgRNA sequences and gene targets for Humagne C+D (enCas12a Library)**

**Table S10: Genome library sgRNA sequences and gene targets for Brunello (Cas9 Library)**

**Table S11: Counts matrix of RNA-sequencing data from unstimulated or IFNγ stimulated human macrophages**
